# Assessment of Protein Complex Predictions in CASP16: Are we making progress?

**DOI:** 10.1101/2025.05.29.656875

**Authors:** Jing Zhang, Rongqing Yuan, Andriy Kryshtafovych, Rachael C. Kretsch, R. Dustin Schaeffer, Jian Zhou, Rhiju Das, Nick V. Grishin, Qian Cong

**Affiliations:** Eugene McDermott Center for Human Growth and Development, University of Texas Southwestern Medical Center, Dallas, TX, USA; Department of Biophysics, University of Texas Southwestern Medical Center, Dallas, TX, USA; Lyda Hill Department of Bioinformatics, University of Texas Southwestern Medical Center, Dallas, TX, USA; Genome Center, University of California, Davis, California, USA; Biophysics Program, Stanford University School of Medicine, Stanford, California, USA; Department of Biochemistry, Stanford University School of Medicine, Stanford, California, USA; Howard Hughes Medical Institute, Stanford University, Stanford, California, USA; Department of Biochemistry, University of Texas Southwestern Medical Center, Dallas, TX, USA; Harold C. Simmons Comprehensive Cancer Center, University of Texas Southwestern Medical Center, Dallas, TX, USA

**Keywords:** CASP16, oligomer prediction, antigen-antibody interaction, stoichiometry, model sampling, AlphaFold2, AlphaFold3

## Abstract

The assessment of oligomer targets in the Critical Assessment of Structure Prediction Round 16 (CASP16) suggests that complex structure prediction remains an unsolved challenge. More than 30% of targets, particularly antibody–antigen targets, were highly challenging, with each group correctly predicting structures for only about a quarter of such targets. Most CASP16 groups relied on AlphaFold-Multimer (AFM) or AlphaFold3 (AF3) as their core modeling engines. By optimizing input MSAs, refining modeling constructs (using partial rather than full sequences), and employing massive model sampling and selection, top-performing groups were able to significantly outperform the default AFM/AF3 predictions. CASP16 also introduced two additional challenges: Phase 0, which required predictions without stoichiometry information, and Phase 2, which provided participants with thousands of models generated by MassiveFold (MF) to enable large-scale sampling for resource-limited groups. Across all phases, the MULTICOM series and Kiharalab emerged as top performers based on the quality of their best models per target. However, these groups did not have a strong advantage in model ranking, and thus their lead over other teams, such as Yang-Multimer and kozakovvajda, was less pronounced when evaluating only the first submitted models. Compared to CASP15, CASP16 showed moderate overall improvement, likely driven by the release of AF3 and the extensive model sampling employed by top groups. Several notable trends highlight key frontiers for future development. First, the kozakovvajda group significantly outperformed others on antibody-antigen targets, achieving over a 60% success rate without relying on AFM or AF3 as their primary modeling framework, suggesting that alternative approaches may offer promising solutions for these difficult targets. Second, model ranking and selection continue to be major bottlenecks. The PEZYFoldings group demonstrated a notable advantage in selecting their best models as first models, suggesting that their pipeline for model ranking may offer important insights for the field. Finally, the Phase 0 experiment indicated reasonable success in stoichiometry prediction; however, stoichiometry prediction remains challenging for high-order assemblies and targets that differ from available homologous templates. Overall, CASP16 demonstrated steady progress in multimer prediction while emphasizing the urgent need for more effective model ranking strategies, improved stoichiometry prediction, and the development of new modeling methods that extend beyond the current AF-based paradigm.

## Introduction

Deep learning techniques have revolutionized the field of protein structure prediction. The field witnessed remarkable progress since CASP13 [1], followed by a breakthrough during CASP14 [2], when AlphaFold2 (AF2) [3] achieved near-atomic accuracy with GDT_TS scores exceeding 90 for most monomer prediction targets. The unprecedented performance of AF2 has transformed biomedical research, and it has become an essential tool for every scientist. Originally designed for predicting monomer structures, AF2 also showed promising performance in protein assembly prediction. DeepMind released AlphaFold-Multimer (AFM) [4] with a similar architecture to AF2 but re-trained on protein assemblies. AFM enabled an unprecedented improvement in protein complex modeling in CASP15 [5]: 90% (37 out of 41) targets had at least one successful model (ICS [6], IPS [6], TM [7], and lDDT [8] all above 0.5) in comparison with 31% in CASP14 when AFM was not yet available to the community [9]. The majority of groups participating in the oligomer prediction category of CASP15 integrated AFM into their pipelines and achieved better performance than the default AFM.

The remarkable progress in CASP15 not only demonstrated how researchers in our community continually advance the frontiers of AI-based protein assembly modeling in the post-AlphaFold era, but also served as a catalyst for innovations in the CASP16 experiment. CASP16 introduced the following novel challenges to stimulate further progress:

**1. Phase 0.** Unlike the standard Phase 1, where stoichiometry is provided, in this phase, participants predict protein complex structures without knowing the stoichiometry. Stoichiometry information is not easy to obtain through experimental methods without solving the 3D structures, and thus, the ability to predict both stoichiometry and 3D structures will make a pipeline more powerful in gaining needed biological insights about protein complexes without the need for complicated experimental assays.
**2. Phase 2.** In Phase 2, the predictors are provided with a large pool of predicted structures— typically 8,040 per target—generated by MassiveFold (MF) [10]. This approach is inspired by a successful strategy used in CASP15, where groups like the Wallner lab [11] demonstrated exceptional performance using extensive sampling approaches such as AFsample [12]. By providing each group with a large set of models, CASP16 aims at accelerating progress in model selection and quality assessment methods.
**3. Model 6.** CASP16 also introduced a "Model 6" submission category alongside the traditional Models 1-5. For Model 6, all participants were required to use the multiple sequence alignments (MSAs) generated by ColabFold [13] (https://casp16.colabfold.com/colabfold_baseline). This experiment is designed to study the influence of MSA quality—a key performance differentiator in CASP15—on prediction accuracy, when separated from other methodological advances. .

By developing AF2 and AFM, DeepMind has played an essential role in driving the progress. After CASP15 and before CASP16, DeepMind developed AlphaFold3 (AF3) [14]. The major innovation of AF3 is its ability to model molecules other than proteins, including DNA, RNA, and small molecules. AF3 also has the potential to set a new standard for protein oligomer prediction. First, due to its reduced dependence on MSAs, AF3 might be more suitable for modeling protein complexes without deep MSAs, such as the interaction between antibodies and antigens. Second, AF3 included a more sophisticated confidence module, and it might have a higher ability in quality assessment and model selection.

Unfortunately, DeepMind did not directly participate in CASP16, but they released AF3 via a web server at the beginning of the experiment, and models from this server were submitted to CASP16 as group AF3-server (run by Arne Elofsson). As a result, many participants used not only AFM but also AF3 in their modeling engines. Meanwhile, similar to CASP15, top-performing groups incorporated multiple strategies to enhance model accuracy, including customized MSAs and extensive model sampling. The winning team for the monomer modeling category, the Yang lab [cite: Yang lab paper], also refined the "constructs" (segments of full sequences) used for modeling, which proved to be another successful strategy for improving model accuracy with AF2/3. We evaluated group performances across all three phases of CASP16, and groups from the Cheng lab [cite: Cheng lab paper] and Kihara lab [cite: Kihara lab paper] appeared to be the overall winners in the oligomer modeling category.

With the release of AF3 and community efforts in optimizing AFM and AF3 performance, we observed a modest improvement in oligomer prediction accuracy in CASP16 compared to CASP15. In addition, antibody/nanobody–antigen (AA) targets presented a pleasant surprise: the winning group, kozakovvajda [cite: Kozakov lab paper], achieved the best performance without using AFM or AF3 to directly model the complexes. By employing a traditional protein– protein docking approach coupled with extensive sampling, they remarkably outperformed both AF3 (with extensive sampling) and the CASP15 winners on AA targets. Due to the limited number of AA targets, we cannot firmly conclude that the kozakovvajda group has made a definitive breakthrough, but we are optimistic about their approach and hope it will inspire the community to explore alternative methods rather than relying exclusively on AF-based strategies. Meanwhile, model accuracy estimation and ranking remain significant challenges. Even the group with the best model-ranking performance was only able to identify their best models for around 60% targets. Finally, we observed a notable improvement in the first model quality in Phase 1 compared to Phase 0, highlighting that correct stoichiometry plays an important role in model accuracy and remains a major challenge in the quest to accurately predict protein structures solely from sequence information.

## Results and Discussion

### Overview of the CASP16 oligomer targets and prediction strategies

The CASP16 oligomer prediction category includes 40 targets (supplemental **Table S1** and supplemental **Fig. S1**) in Phase 1 (the main experiment), among which 22 are hetero-oligomers and 18 are homo-oligomers. 30 of these targets were used in Phase 0 to challenge predictors’ ability to predict stoichiometry, and 35 of them were included in Phase 2. More than half (21 of 40) of the target structures were determined by cryogenic electron microscopy (cryo-EM), and the rest were solved by X-ray crystallography.

Most of these target complexes are formed by proteins from the same organism, whether cellular (bacterial or eukaryotic) or viral. We expect the coevolutionary signals between components of these complexes can be used to guide complex prediction. Two targets represent host-pathogen interactions. Predicting protein-protein interactions (PPIs) across species is more challenging because cross-species interactions tend to have a shorter evolutionary history, limiting the coevolutionary signals [15–17]. However, the performance on these two targets is moderate (95% quantile of DockQ [18] for these targets is about 0.5 and 0.6, respectively), possibly related to the fact that pathogen proteins were targeting surfaces known to mediate PPIs. Finally, there are three nanobody-antigen and five AA complexes. Due to the lack of coevolutionary signals in such complexes, they are expected to be the most challenging [19] and are a focus of our evaluation.

Although predictors are supposed to predict structures with known stoichiometry in Phase 1, there are two filament targets in Phase 1 whose stoichiometry cannot be defined, i.e., T1219o and T1269o. T1219o is a homo oligomer composed of human alpha-defensin-6 solved by cryo-EM. The structure (1zmq) of human alpha-defensin-6 was determined two decades ago [20], but this target represents a higher-order oligomeric complex than the known experimental structure (supplemental **Fig. S2**), where negatively charged phosphates were trapped in between different subunits and are possibly mediating their interactions. The predictors were not aware of the presence of phosphates, and none of them were able to predict the conformation as in the target, but several groups might have utilized the structure of the asymmetric unit in 1zmq for the prediction. Similarly, T1269o is also a filament target with multiple PDB templates (7a5o [21] and 8oes) with an identity above 50% and nearly full target coverage. Finally, for target T1295o, while the predictors were asked to predict based on a stoichiometry of A8, the biological assembly of this target consists of 24 chains (complex size beyond what most groups can accommodate). Therefore, we treated T1295o similarly as Phase 0 targets due to the difference in stoichiometry between the target and the models.

In this CASP, 13,630 models were submitted by 87 groups in Phase 1, which is the same oligomer prediction challenge as in CASP15. This marks a 20% increase in the number of submitted models compared to CASP15, despite no change in the number of participating groups. An overview of the performance of each group on each target measured by best DockQ is shown in **Fig. 1A**, and only groups that submitted models for more than 10 targets (25%) in Phase 1 are included. In contrast to what we observed in the monomer category [cite: our monomer evaluation paper], where most groups can generate high-quality models for most targets, predictors’ performance in the oligomer category is much worse. About 50% of targets cannot be correctly predicted by most groups (DockQ < 0.5, white to blue cells in **Fig. 1A**). For most of the more challenging targets, there are several groups that were able to generate at least medium-quality models (DockQ >= 0.5, white to red cells in **Fig. 1A**), but not a single group can consistently generate such models across multiple challenging targets (supplemental **Fig. S3**).

**Figure 1.**
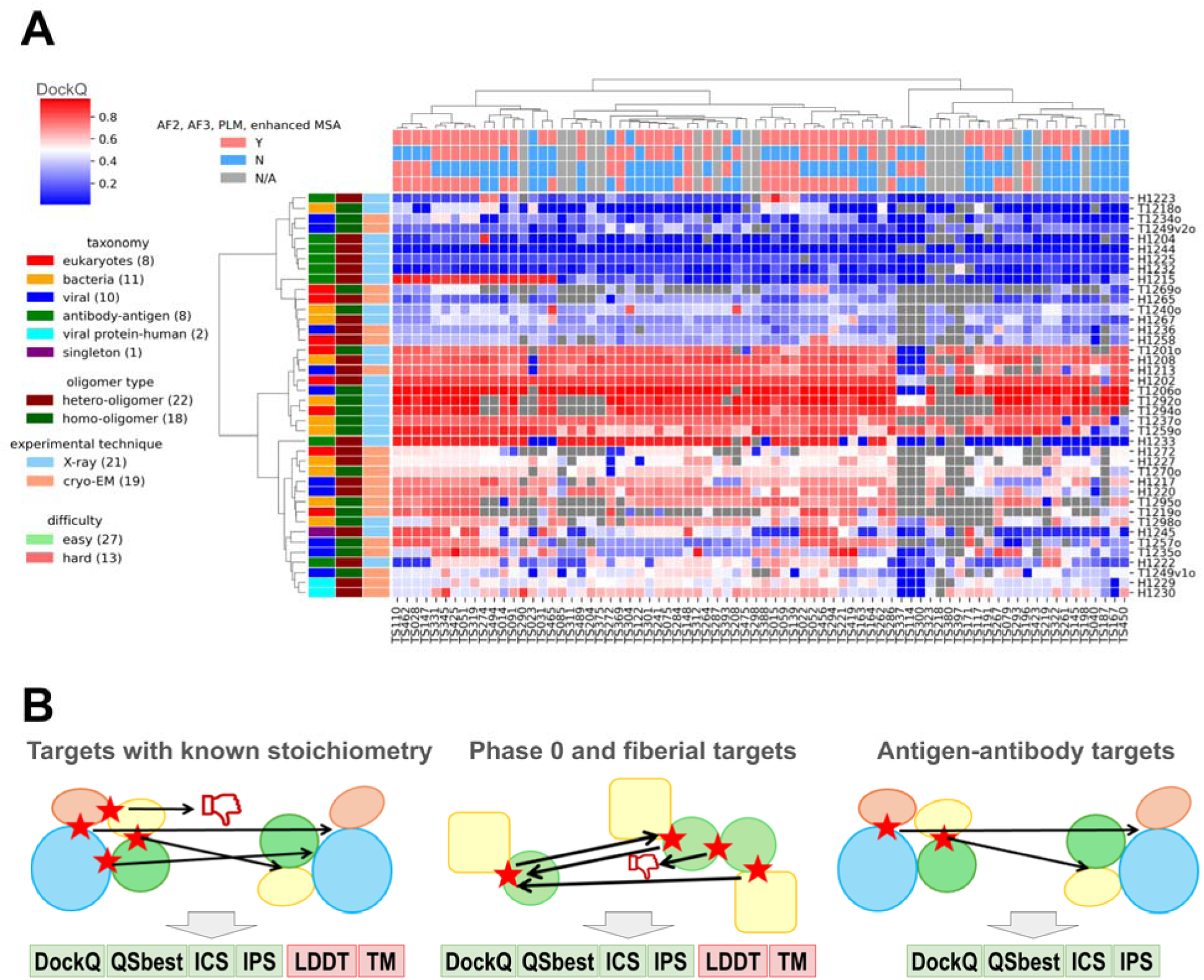
Overview of targets and participating groups. **A)** A heatmap of the performance of participating groups over Phase 1 oligomer targets. The value in each cell is the highest DockQ value of one group for one target (Grey indicates that such a score is not available). The annotations on the x-axis and y-axis represent the different features of groups and targets, respectively. Only groups that submitted more than 25% of the models are included. **B)** Pipelines we used to evaluate performance for different categories of targets. Left: a standard pipeline for most Phase 1 and Phase 2 targets, where targets share the same stoichiometry as the models. Middle: a special pipeline for Phase 0 and special targets in other phases, where models have different stoichiometry from the targets. Right: a pipeline for AA targets focusing on antibody–antigen interfaces.

Similar to what we did in CASP16 monomer evaluation, we annotated each group by the strategies they used (**Fig. 1A**). The majority of groups used either AFM or AF3 or both. Consistent with monomer prediction, the groups incorporating AF3 results have a noticeable improvement over groups using only AF2 (supplemental **Fig. S4**). Nearly all groups providing abstracts used MSA. About half of them tried to enhance MSAs over the AFM’s MSA generation pipeline, and their performance was better than groups without using enhanced MSAs (supplemental **Fig. S4**). In addition, in this CASP, there is no group directly using protein language models (PLM) for structure prediction, instead, some groups incorporated the encodings from PLMs such as ESM2 [22] for sequence searches or MSA construction. Groups that used PLM show worse performance than those that did not (supplemental **Fig. S4**). However, we believe that this is not due to PLMs having a negative impact; instead, it might be related to that groups who used PLM did not engineer the other aspects of their modeling pipeline as diligently as other groups.

### Overview of our evaluation strategies

The additional challenges (Phase 0) and special targets described above required specialized evaluation routines. The default strategy for oligomer target evaluation, implemented at the CASP Prediction Center, utilizes the OpenStructure (OST) software [23,24] to compute a number of metrics to evaluate the agreement between target PPI interfaces and model PPI interfaces, including ICS [6], IPS [6], DockQ [18], and QSbest [25], among others. OST also computes metrics assessing overall structural similarity between targets and models, such as TM-score [7] and lDDT [8]. To calculate these scores, OST first establishes a one-to-one mapping between chains in the target and chains in the model. Mapping initially relies on sequence similarity, attempting to match each target chain to its most similar chain(s) in the model. In cases of multiple-to-multiple matches, OST selects the best one-to-one mapping that maximizes structural similarity between the target and the model (**Fig. 1B** left).

While the OST pipeline is suitable for most Phase 1 targets, where the target and model share the same stoichiometry, it is not appropriate for Phase 0 and certain Phase 1 targets (T1219o, T1269o, and T1295o) where stoichiometry differences exist between the target and the model. When a model contains more chains than the target, OST maps the best matches for each target chain to maximize similarity, ignoring unmapped chains—even if they contain incorrectly predicted interfaces (supplemental **Fig. S5**). Conversely, when a model contains fewer chains than the target, even if all critical interfaces are correctly modeled, the model is heavily penalized (supplemental **Fig. S5**). Consequently, the OST-based evaluation may favor models predicting excess copies of each subunit over models with the correct stoichiometry. In extreme cases, for a target with A1B1 stoichiometry, one could artificially inflate scores by including multiple copies of A1B1 complexes within a single model and relying on OST to select the best match.

To address this limitation, we introduced an alternative evaluation pipeline termed Reciprocal Best Match (RBM) for targets where stoichiometry between the target and the model could differ. In the RBM pipeline, we compare target and model structures at the level of interacting chain pairs (**Fig. 1B** middle). Specifically, for each pair of interacting chains in the target (defined by minimal inter-residue distance <5L), we search for the best matching pair in the model to maximize each similarity score (ICS, IPS, DockQ, QSbest, TM-score, and lDDT). This approach resembles the CAPRI evaluation scheme, although we used additional scores beyond those incorporated in DockQ [26,27]. Importantly, we also initiated matching from each interacting chain in the model, seeking the best corresponding pair in the target. The final evaluation was based on the weighted average (weighted by interface size; see **Methods**) of the target-to-model and model-to-target scores for each metric. This dual scoring assesses both how well the model captures target interfaces and whether all predicted model interfaces are present in the target. We applied this RBM pipeline to all Phase 0 targets and to three Phase 1 targets (T1219o, T1269o, and T1295o). We introduced the RBM pipeline during CASP16 evaluation, and we hope it will establish a foundation for future assessments. Phase 0, which examines predictors’ ability to model complex structures solely from sequence, should become a main focus of future CASP experiments, and we urge future assessors to carefully consider appropriate evaluation metrics for this challenge.

Finally, both the OST and RBM pipelines have limitations when applied to AA targets. In these cases, the only critical interface is the interface between the antibody and its antigen. However, many CASP16 AA targets included multiple antibody and antigen chains, and trivial interfaces between antibody–antibody or antigen–antigen chains were often large and easily predicted. These trivial interfaces could dominate OST and RBM scoring, artificially inflating scores and masking failures to accurately model the critical interface between antigens and antibodies. To address this, we classified chains into antibody/nanobody and antigen groups and evaluated only interfaces between these two classes. We also excluded TM-score and lDDT from the evaluation of AA targets, as the relative global orientation of an antibody and antigen is less important than accurately modeling their interface (**Fig. 1B** right). This additional evaluation procedure, built on top of the OST pipeline for Phase 1 and Phase 2 AA targets and the RBM pipeline for Phase 0 AA targets, is referred to as the AA pipeline.

### Features of challenging oligomer targets in CASP16

We assessed model quality across CASP16 targets using the 95th percentile of DockQ and TM-scores, as shown in **Fig. 2A**. Targets with DockQ < 0.4 or TM-score < 0.8 were classified as difficult targets (easy targets inside the green box in **Fig. 2A**). Five out of the eight AA targets are regarded as difficult (red dots in **Fig. 2A**). Other difficult targets, shown in **Fig. 2B – 2I**, include examples with low interface accuracy (e.g., H1234 and H1236 with low DockQ scores) or incorrect global topology (e.g., T1269o and H1265 with low TM-scores). Frequently, the incorrect global topology is caused by mispredicted or missing interfaces. When these interfaces are small, they do not contribute significantly to the interface-based scores, but they might change the shape of the complex and are thus penalized by TM-scores. Below, we discuss our observations about some of these difficult targets and exceptional models for them.

**Figure 2.**
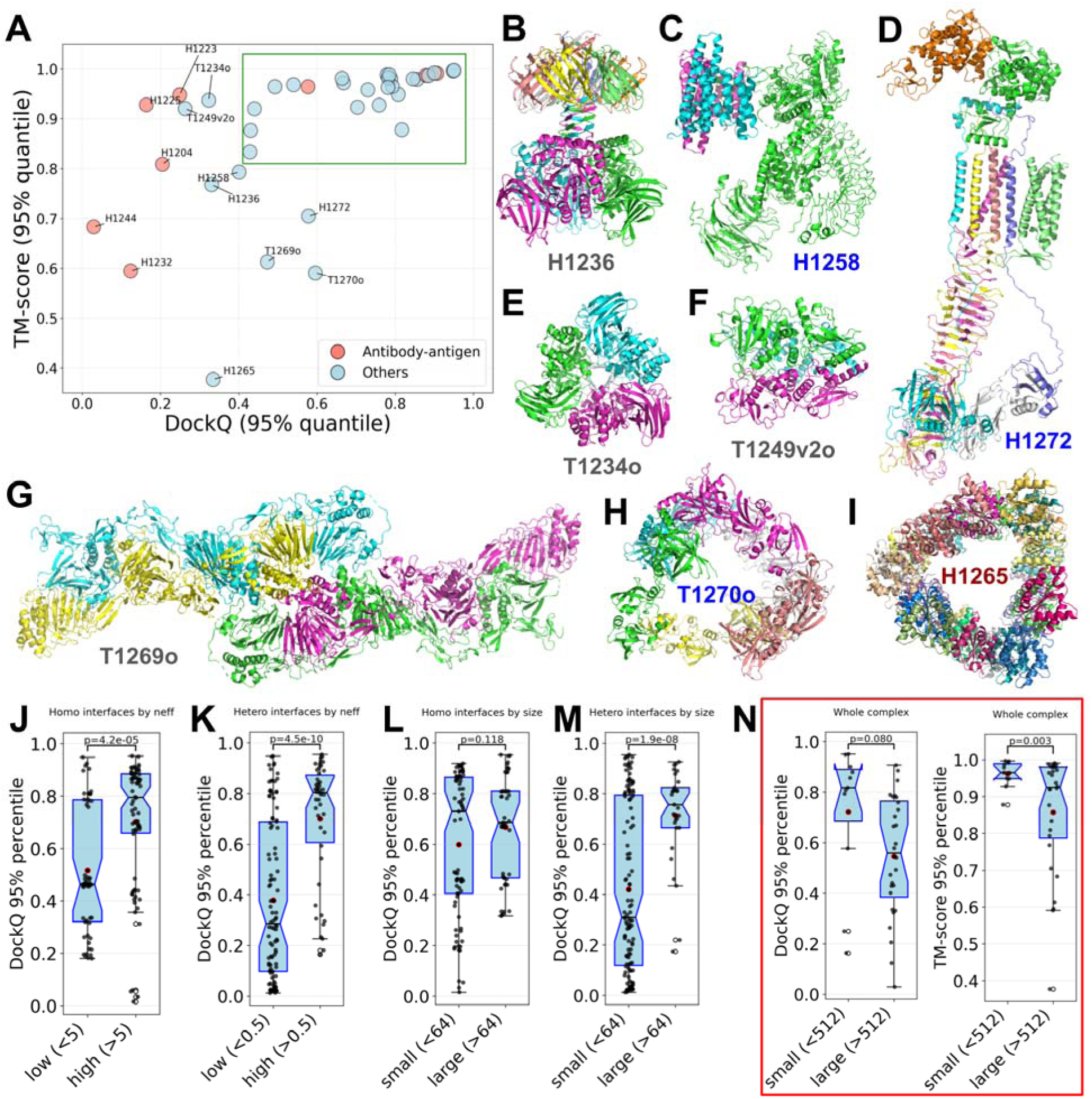
Difficult targets and target properties that affect model quality. **A)** Identification of difficult oligomer targets in CASP16 using the 95% quantile of DockQ and TM-score. Targets inside the green box are considered easy. **B – I)** Structures of identified difficult targets. Each chain is indicated by a different color. **J – N)** Target features affecting model quality: **J)** *Neff* of input MSAs for homo-dimeric interfaces (each dot represents an interface); **K)** *Neff* of input MSAs for hetero-dimeric interfaces (each dot represents an interface); **L)** interface size (the number of inter-chain contacts with distance below 5L) for homo-dimeric interfaces (each dot represents an interface); **M)** interface size (the number of inter-chain contacts with distance below 5L) for hetero-dimeric interfaces (each dot represents an interface); **N)** Total number of residues in the entire complex (each dot represents a target).

H1236 (**Fig. 2B**) comes from Haloferax tailed virus 1, and it represents a complex formed by the viral capsid protein Gp30 and the prokaryotic polysaccharide deacetylase. Most top-performing groups predicted the overall structure of this target reasonably well. However, only KiharaLab/kiharalab_server [cite: Kihara’s paper] accurately modeled the interface, achieving a DockQ above 0.6. Compared to other groups, KiharaLab uniquely captured the N-terminal intertwined β-sheet formed by the first eight residues of six Gp30 subunits (top of **Fig. 2B**), contributing to their superior performance. However, similar to other groups, they did not predict the additional intertwined β-sheets (middle of **Fig. 2B**) formed by the first 35 N-terminal residues of the prokaryotic polysaccharide deacetylase (supplemental **Fig. S6**).

H1258 (**Fig. 2C**) is a complex between human LRRK2 (Leucine-Rich Repeat Kinase 2) and its regulatory protein 14-3-3 [28]. Two groups from the Yang lab [cite: Yang’s paper] achieved DockQ scores above 0.7, remarkably outperforming other groups in interface quality. Notably, instead of modeling full LRRK2 in the complex, they only modeled residues 861–1014 of LRRK2, the domain interacting with 14-3-3 in the target. Therefore, although they have superior DockQ, their TM-score and lDDT are lower than average. The Yang lab used an innovative strategy of iterative modeling by refining the constructs (segments of full protein sequences) to be used as AF2/3 inputs. In this case, modeling with only the 14-3-3 interacting domains likely helped them to obtain a more precise LRRK2/14-3-3 interface.

Another difficult oligomeric target, T1249v2o (**Fig. 2F**) is a homo-oligomer formed by an Arenaviral spike protein. Homo-oligomeric complexes of this protein adopt two noticeably different conformations, open (T1249v1o) and closed (T1249v2o) states, based on experimental data. Both states were used as CASP16 targets, and the predictors were challenged with the task of predicting two possible conformations. However, predictions for both targets mostly resemble the open state, leaving the closed state (T1249v2o) a challenging target and highlighting the difficulty of predicting multiple conformations.

Another difficult case, T1270o, is a hexamer of HtrA protease from *Borreliella burgdorferi,* the pathogen causing Lyme disease (**Fig. 2H**). While many models achieved good DockQ scores of around 0.6, their TM-scores are close to 0.5, indicating a poor agreement of overall shape between the target and its models. The target is a hexamer, organized as two trimmers, each forming a triangular arrangement through interactions among their N-terminal domains. These two trimers are connected via three additional interfaces involving only the C-terminal domains. Detailed examination of models with high DockQ scores (≥ 0.6) revealed that, although the N-terminal interactions within each trimer were predicted with moderate accuracy, most models failed to accurately capture the C-terminal interfaces between trimmers. One striking feature of this target is that this complex is highly not compact, leaving a large empty hole in the middle of the structure. However, AFM and AF3 tend to predict compact structures and models submitted by the predictors also mostly failed to predict the large empty space in the middle. This large space in the middle is possibly important for HtrA’s protease function, allowing it to trap its substrate inside the complex; nevertheless, the lack of compactness for such targets poses challenges to the modern structure prediction methods.

Finally, a complex with the highest number of chains among CASP16 targets, H1265 (**Fig. 2I**), turned out to be particularly challenging for all CASP16 groups. H1265 is a complex of Toll/Interleukin-1 Receptor (TIR) domains of the Toll-like receptor 4 (TLR4), which plays a crucial role in innate immune signaling [29]. Not a single group correctly predicted the topology of this target, and it shows the lowest TM-score among all targets. The best model from the Kihara lab captured the overall triangular shape of the target. But they predicted interfaces correctly, leading to a different handedness from the target structure. Other group’s models exhibit shapes totally different from that of the target (supplemental **Fig. S7**). H1265 consists of two types of sequences with a stoichiometry of A9B18, and in order to form the whole complex, the two sequentially different subunits interact with each other in four possible ways. Based on our own experience, if two proteins, A and B, have multiple possible interfaces, their interactions are harder to predict due to the difficulty of disentangling the coevolutionary signals arising from different interfaces [30]. With multiple possible interfaces between A and B and an unusual triangular shape, it is not surprising that H1265 turns out to be a particularly challenging target.

In parallel to manual inspection of difficult targets, we also examined general factors influencing prediction accuracy. We performed these analyses for all non-redundant pairwise PPI interfaces in the oligomeric targets. Each oligomer target may contribute multiple interfaces, improving the statistical power of our analyses. We also partitioned these binary PPI interfaces into homo-dimeric and hetero-dimeric. Consistent with findings from previous CASP experiments, the diversity of sequences in the MSAs or paired MSAs (for hetero-dimeric interfaces), as measured by *Neff* [31], significantly affects the modeling accuracy of homo-dimeric (**Fig. 2J**) and hetero-dimeric interfaces (**Fig. 2K**). *Neff* appears to have a stronger impact on the quality of hetero-dimeric interface predictions compared to homo-dimeric ones. Another known factor influencing complex prediction accuracy is the interface size [32]. We found that interface size has little effect on the prediction accuracy of homo-dimeric interfaces (**Fig. 2L**) but is more influential for hetero-dimeric interfaces (**Fig. 2M**), where smaller interfaces are generally harder to predict. Several oligomer targets are huge in size and might be challenging to fit into GPUs. The number of residues in CASP16 oligomer targets ranges from 300 to around 6000. We wondered if the size of a complex would affect prediction accuracy, and indeed found that larger complexes tend to show poorer performance based on both DockQ and TM-score (**Fig. 2N**).

### A moderate progress in oligomer prediction

To evaluate whether the field has made progress from CASP15 to CASP16, we compared predictor performance across the two experiments. Because the apparent performance in different CASP rounds can be significantly influenced by the overall difficulty of the targets, we needed a method that remained consistent between CASP15 and CASP16 to serve as an indicator of target difficulty. We used the performance of ColabFold [13] as this indicator. ColabFold participated in CASP15 and submitted predictions for all targets. For CASP16, the similar ColabFold pipeline with updated databases and AFM weights (from multimer-v2 to multimer-v3), was applied to 37 out of 40 Phase 1 targets to help estimate target difficulty. Filamentous targets without defined stoichiometry and very large targets that could not be accommodated in GPU memory were excluded from ColabFold modeling. Accordingly, our comparative analysis was restricted to targets for which ColabFold predictions were available.

We partitioned the CASP15 and CASP16 targets with ColabFold predictions into normal (non-AA) and AA targets, and first focused on the normal targets. We calculated the average DockQ scores of the first models from the top 55 groups in CASP15 and CASP16 (**Fig. 3A**). The highest DockQ among CASP participants increased from 0.55 to 0.59. However, the DockQ score of ColabFold also increased—from 0.43 in CASP15 to 0.47 in CASP16—suggesting a potential shift in the overall difficulty of the targets. The distribution of DockQ scores across targets further supports this interpretation, with CASP15 containing a higher proportion of difficult non-AA targets compared to CASP16 (**Fig. 3B**). The closeness of these numbers suggests that there has not yet been another major breakthrough in oligomer structure prediction, although whether measurable progress has been made requires more robust statistical analysis. One notable difference is that a greater number and fraction of groups clearly outperformed ColabFold in CASP16, indicating that the field is beginning to surpass AF2, possibly due to the contributions of AF3.

**Figure 3.**
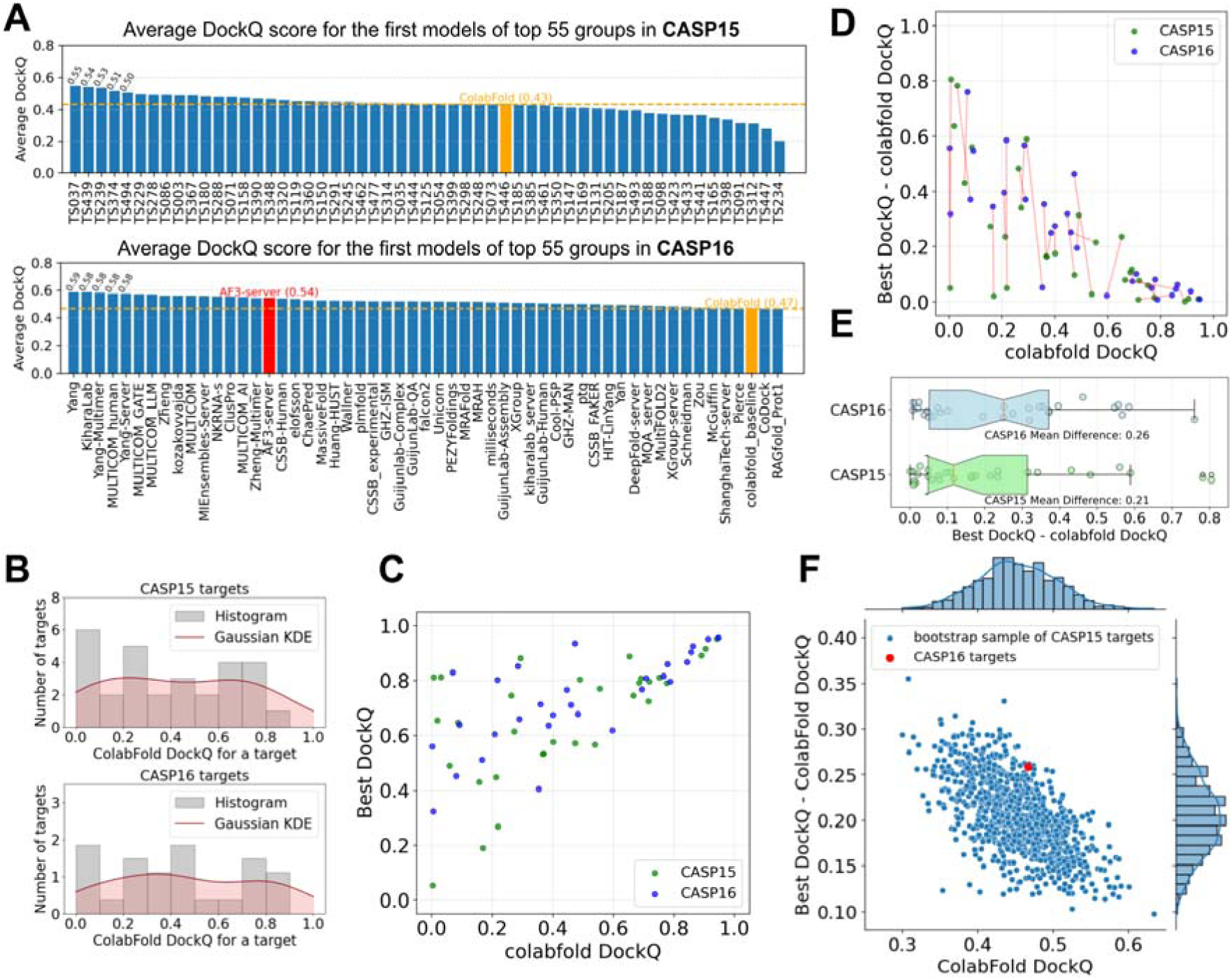
Moderate progress in CASP16 over CASP15 after accounting for differences in target difficulty. **A)** Average DockQ for top 55 groups over normal targets (excluding AA and fiber targets) in CASP15 (top) and CASP16 (bottom), respectively. **B)** Gaussian kernel density estimation of the DockQ distribution of the first model of ColabFold for CASP15 (top) and CASP16 (bottom) targets. **C)** The quality of the best models and the ColabFold models, measured by DockQ. **D)** One-to-one mapping between CASP15 and CASP16 targets by the Hungarian algorithm to sample targets from CASP15 to match the target difficulty level of CASP16 targets. **E)** Difference in DockQ between the best models and the ColabFold models for CASP16 targets and CASP15 targets sampled by the Hungarian algorithm, respectively. **F)** Performance comparisons between CASP16 targets and weighted bootstrap samples of CASP15 targets. These bootstrap samples of CASP15 targets show similar difficulty levels as CASP16 targets (by ColabFold DockQ, x-axis) and improvement over the ColabFold models by the best CASP15 or CASP16 models (y-axis).

To evaluate progress more robustly, we first examined the relationship between target difficulty, as indicated by the DockQ score of ColabFold’s first model, and the field’s optimal performance (in CASP15 or CASP16) on each target (**Fig. 3C**). Differences in target difficulty can complicate evaluation of progress: easier targets offer limited room for improvement over the ColabFold baseline, whereas harder targets correlate with larger potential performance gains between the best model and the ColabFold model. To more objectively assess progress between CASP15 and CASP16, we implemented two statistical approaches designed to mitigate biases arising from variation in target difficulty.

First, we applied the Hungarian algorithm [33], which finds the optimal one-to-one matching between two sets, to select a subset of CASP15 targets that closely matched the difficulty level of CASP16 targets. In our case, the set of matched targets (connected by red lines in **Fig. 3D**) showed very similar overall difficulty levels between the two CASPs, with average ColabFold DockQ scores of 0.456 for CASP15 and 0.467 for CASP16. Based on this matching to control for shifts in target difficulty, we calculated the performance improvement over ColabFold (defined as the difference between the best DockQ and ColabFold’s DockQ) for CASP15 and CASP16 (**Fig. 3E**). We found that in CASP16 (mean improvement: 0.26), the community tended to achieve noticeably better performance than in CASP15 (mean improvement: 0.21), although the difference was not statistically significant (paired t-test p-value = 0.184) due to the limited number of targets.

Second, we used a weighted bootstrap technique to generate samples (with replacement) of CASP15 targets, aiming to match the ColabFold DockQ distribution of CASP16 targets. For each bootstrap sample of CASP15 targets, we computed the average ColabFold DockQ (indicating target difficulty) and the average improvement of the best model’s DockQ over ColabFold’s DockQ (performance improvement over baseline), with the distributions of these two measurements shown in **Fig. 3F**. Across these bootstrap samples, we observed a clear negative correlation between overall target difficulty and performance improvement. Among the CASP15 bootstrap samples with difficulty levels similar to CASP16 (near the red dot representing CASP16), very few achieved better overall performance than CASP16. This further supports the conclusion that measurable progress has been made in oligomer prediction accuracy. However, we speculate that much of this moderate improvement is attributable to the introduction of AF3, a new strategy available to CASP16 participants.

### In-depth study of antibody/nanobody-antigen targets

AA complexes have attracted significant attention in CASP experiments due to their biological and therapeutic importance, yet accurate modeling of these complexes remains a major unsolved challenge. One primary advantage of AF3 over AF2 is its improved ability to model AA complexes. CASP16 included eight AA targets in Phase 1, matching the number of such targets in CASP15. To assess progress, we compared the average DockQ scores of the top 68 participating groups across the two CASPs (**Fig. 4A**). The highest average DockQ in CASP15 was 0.5, which slightly decreased to 0.47 in CASP16. At first glance, this may suggest stagnation or even a decline in performance. However, it is important to note that ColabFold— used as a consistent baseline—achieved an average DockQ of 0.21 in CASP15 but dropped sharply to 0.05 in CASP16, indicating that the AA targets in CASP16 were remarkably more difficult. It is important to note that the majority of groups in CASP16 were able to outperform ColabFold in AA targets, indicating an improvement in the field.

**Figure 4.**
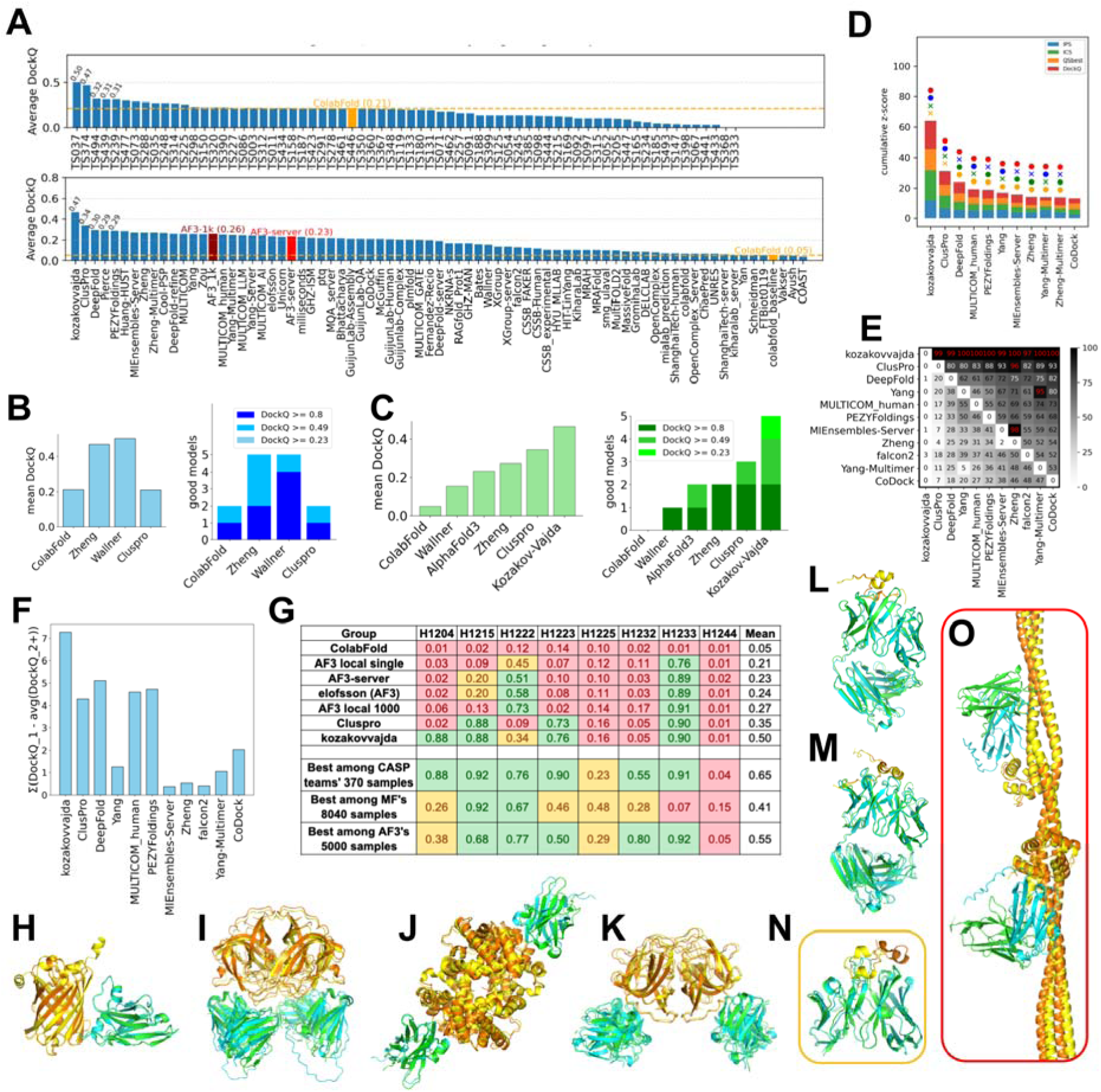
Performance evaluation on AA targets. **A)** Average DockQ scores of first models for AA targets submitted in CASP16 (bottom) and CASP15 (top) by the top 66 groups. Dashed orange lines indicate the DockQ of ColabFold in both CASPs. We added AF3_1k, which uses the best model of each target selected by ipTM of the antibody–antigen interfaces among 5,000 models we generated using 1,000 random seeds with AF3. **B)** Left: average DockQ of first models from selected groups in CASP15. Right: number of targets with good models falling into different quality tiers based on DockQ for selected CASP15 groups. **C)** Left: average DockQ of first models from selected CASP16 groups. Right: number of targets with good models falling into different quality tiers based on DockQ for selected CASP16 groups. **D)** Ranking of groups based on their cumulative z-scores (z-score for each target is the average of 4 component scores, ICS, IPS, QSbest, and DockQ) for AA targets in all phases. **E)** Pairwise head-to-head bootstrap comparisons between groups for AA targets in all phases. Each cell shows the percentage of targets where the row group outperforms the column group based on cumulative z-scores. **F)** The sum of differences between each group’s first model DockQ and the mean DockQ of all other models per target. **G)** Per-target DockQ of models from representative CASP16 groups or generated by us using a standalone AF3 across CASP16 AA targets (labeled on top). AF3 local single: the average of the first models generated by AF3 using 1,000 random seeds; AF3 local 1000: best model selected by ipTM of antibody–antigen interfaces from 5,000 AF3 models generated using 1,000 random seeds. **H–O)** Structural superimpositions of targets and the best models selected by cumulative DockQ of antibody–antigen interfaces for AA targets. Targets without high-quality models are in orange (**N**: H1225) and red (**O**: H1244) boxes. Orange: antigen chains in targets; yellow: antigen chains in models; green: antibody chains in targets; cyan: antibody chains in models.

This difference in target difficulty should be taken into account when interpreting trends across CASP experiments. However, due to the small number of AA targets, we could not apply the same difficulty-matching strategies used to evaluate the progress in non-AA targets. Fortunately, the top-performing groups in CASP15, Wallner (TS034) [11] and Zheng (TS374) [34], also participated in CASP16, allowing us to use them as a reference for progress evaluation under the assumption that their prediction pipeline did not decline in performance. A review of their method descriptions confirmed that Wallner and Zheng used improved versions of the methods they used in CASP15, i.e., AFsample [12] and DMFold-Multimer [35], respectively.

In CASP15, ClusPro [36] performed worse than Wallner and Zheng and was on par with ColabFold [9], with a mean DockQ around 0.2 and only two acceptable models (DockQ ≥ 0.23), while Wallner and Zheng each predicted five acceptable targets (**Fig. 4B**). In contrast, in CASP16, both ClusPro (server group) and kozakovvajda (human group form the same lab) outperformed Wallner and Zheng: ClusPro and kozakovvajda achieved acceptable accuracy on three and five targets, respectively, while Wallner and Zheng succeeded on only one and two targets, respectively (**Fig. 4C**). ColabFold failed to generate acceptable models for any CASP16 AA targets. Assuming the predicting capacity of methods from Wallner and Zheng did not decline, the superior performance of ClusPro and kozakovvajda suggests genuine methodological advancements in AA target prediction. However, given the small number of AA targets, we lack the statistical power to draw definitive conclusions.

Nevertheless, the kozakovvajda group was the clear winner in the AA modeling category of CASP16, significantly outperforming other groups in the accuracy of first models as supported by head-to-head bootstrapping analysis (**Fig. 4D** and **4E**). Most other top-performing groups did not show statistically significant differences from each other (**Fig. 4E**). In addition, the kozakovvajda group excels at ranking the AA targets, with 5 of their first models being the best and showing the largest score difference between their first model and other models (**Fig. 4F**).

The improved performance in AA interactions is a major advantage of AF3 over AF2. However, we were surprised to observe that many groups outperformed the AF3 server. Considering that most groups employed large-scale model sampling—and that AF3’s reported good performance on AA complexes is tied to large sampling size—this result becomes less unexpected, because the AF3 server group (performed by the Elofsson lab [cite: Arne’s paper]) likely only sampled 5 models for each target. To enable a fairer comparison, we conducted extensive sampling with a standalone version of AF3 (using 1,000 seeds and generating 5,000 models, referred to as AF3_local_1000) and evaluated whether kozakovvajda and ClusPro still maintained superior performance. As shown in **Fig. 4G**, AF3_local_1000 produced only two models with DockQ scores above 0.49 and achieved a mean DockQ of ∼0.27 across the eight targets—still well behind the kozakovvajda group.

We wondered whether the large number of models generated by MF and AF3 contained the correct conformation for every AA target (**Fig. 4G** below). Among the over 8,000 MF models, we were able to find high-quality predictions (DockQ >= 0.49) for two out of eight AA targets. AF3’s sampling seems to be more effective than MF, among which we can find high-quality models for five targets. The ∼370 models submitted by CASP16 groups appear to be the best pool of sampled conformations, containing high-quality models for six targets. These results underscore the strength of community contributions and collaborative innovation in advancing AA complex modeling. Nevertheless, the massive sampling still missed the interface for some AA targets, indicating that effective sampling remains an unsolved problem for AA modeling.

Structures in **Fig. 4H – 4O** present structural superpositions between AA targets and the best predicted models. Most targets (**Fig. 4H – 4M**) have models predicted with high accuracy among CASP16 predictions. One target, H1244 (**Fig. 4O**), is particularly challenging; CASP16 groups, MF, and AF3 all failed to obtain an acceptable model for this target among the massive number of predictions. The antigen in H1244 consists of an EF-hand domain and an extended coiled-coil, and the antibody interacts with this antigen via a small interface (green and orange proteins in **Fig. 4O**) without interacting with the EF-hand domain. However, predicted structures tend to have larger interfaces between the antibody and antigen (cyan and yellow proteins in **Fig. 4O**) and involving the EF-hand domain. Another difficult target was H1225 (**Fig. 4N**), between the CDRs of an antibody and the respiratory syncytial virus attachment glycoprotein, where only acceptable models (max DockQ = 0.48) were obtained among CASP16 participants or massively sampled MF and AF3 models.

### Performance evaluation and ranking

#### Phase 1 ranking

Phase 1 is the main protein oligomer prediction stage of CASP16, continuing the traditional CASP oligomer challenges that began in CASP12 (2016) [6], where both the sequences and stoichiometry are provided. Group performance was evaluated using cumulative z-scores across all Phase 1 targets based on the best model for each group (**Fig. 5A**). The leaderboard was dominated by the MULTICOM series [cite: Jianlin Cheng’s paper], with four of the top five groups being MULTICOM_AI, MULTICOM_human, MULTICOM_LLM, and MULTICOM_GATE. Kiharalab [cite: Kihara’s paper] was the only non-MULTICOM team to break into the top five, followed by other strong performers such as Yang-Multimer [cite: Yang’s paper], and CSSB_FAKER. These top-performing groups generally achieved high overall structure accuracy (TM-score and lDDT), as well as high interface quality (DockQ, ICS, IPS, and QSbest). Interestingly, while most leading groups performed well across multiple metrics, kozakovvajda distinguished itself by excelling primarily in interface quality (DockQ and ICS), but not in overall structure accuracy (TM-score and lDDT). This is because they used a protein-protein docking-based pipeline, different from most other CASP16 groups.

**Figure 5.**
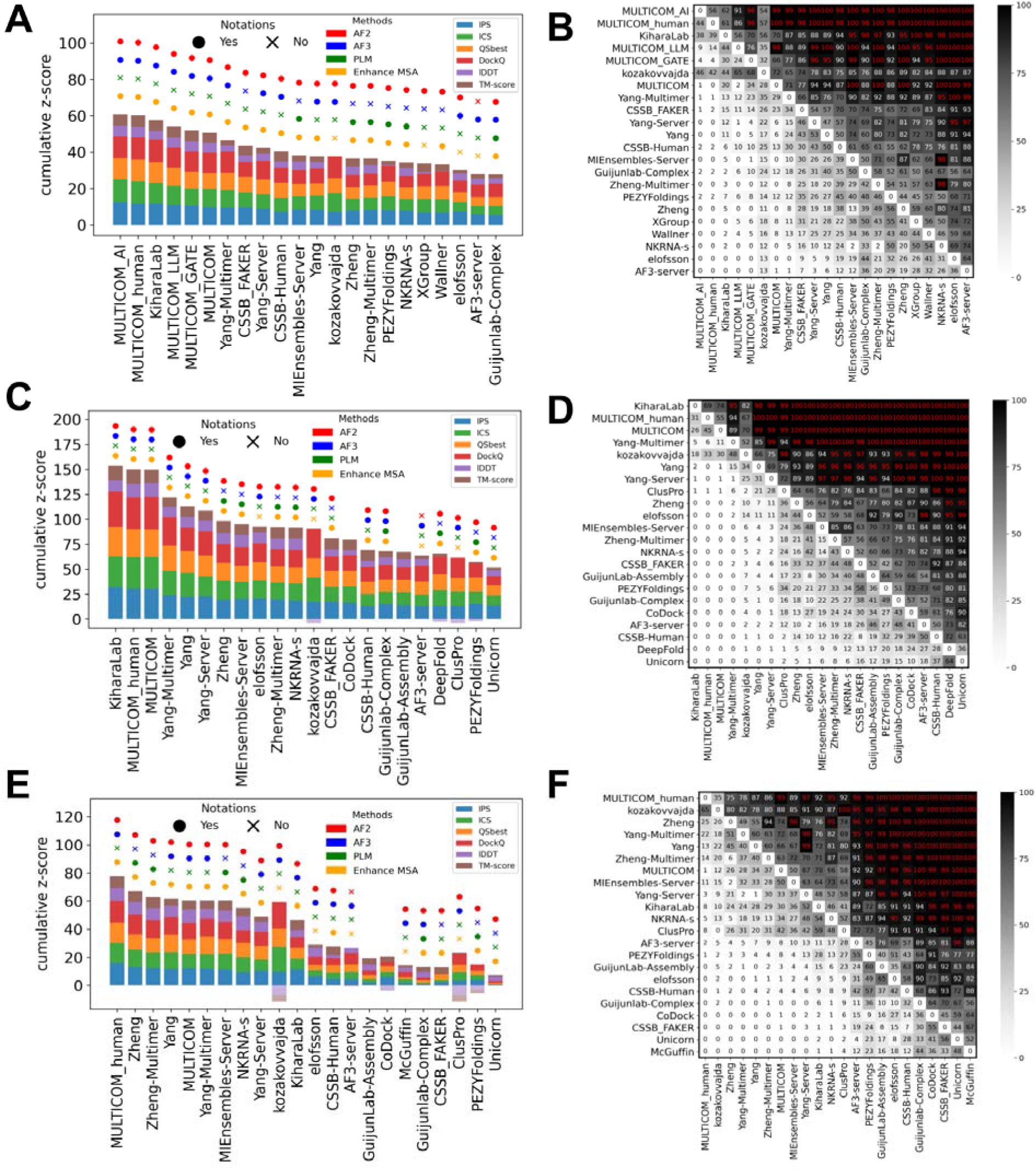
Ranking based on cumulative z-scores and head-to-head comparisons between groups using bootstrap samples. **A)** Ranking by best models for Phase 1 targets. **B)** Head-to-head comparisons by the best models for Phase 1 targets. **C)** Ranking by best models for all targets in three phases. **D)** Head-to-head comparisons by the best models for all targets in three phases. **E)** Ranking by first models for all targets in three phases. **D)** Head-to-head comparisons by first models for all targets in three phases.

To provide a statistical measure of performance difference, we performed head-to-head comparisons with bootstrapping. The results are shown in **Fig. 5B**, where each entry in the matrix reflects the percentage of bootstrap samples in which one group outperformed another based on cumulative z-scores. These statistical tests suggest that there is no single winner. MULTICOM_AI significantly outperformed all non-MULTICOM groups except Kiharalab and kozakovvajda; Kiharalab, however, does not significantly outperform many following groups. This lack of significant differences in performance among top groups is likely attributable to their common reliance on AFM or AF3 as the core prediction engine, combined with similar performance-enhancing strategies such as customized MSAs and extensive model sampling.

We also evaluated group rankings in Phase 1 based on model 1, i.e., the preferred model by each group. While the rankings differed slightly from those based on best models, the MULTICOM series, Yang-Multimer, and Kiharalab remained top performers. However, performance differences between groups became even more marginal, lacking statistically significant differences among the top 20 groups (**Fig. S8A** and **S8B**). This suggests that the top-performing groups in the best models do not have additional advantages in ranking models correctly. Many CASP16 groups use similar scoring strategies, frequently relying on individual or combined AF2/3 confidence metrics such as pLDDT, ipTM, and pTM scores.

#### Ranking over all Phases

Unlike previous CASP experiments, CASP16 introduced two additional phases. To determine an overall ranking across all three phases, we calculated cumulative z-scores and conducted head-to-head comparisons using targets from all phases. To stress the importance of the additional CASP16 challenges, **we consider the ranking over all phases as the main CASP16 ranking**. Rankings were based on both the best (**Fig. 5C** and **5D**) and the first models (**Fig. 5E** and **5F**). When evaluated using the best models, Kiharalab ranked first, overtaking MULTICOM_AI, which had led in Phase 1. MULTICOM_human and MULTICOM followed closely, with very narrow performance gaps among the top three and no significant differences (**Fig. 5D**). Nevertheless, these groups significantly outperformed most others, with the exceptions of kozakovvajda and Yang-Multimer.

Notably, rankings shifted considerably when the first models were used (**Fig. 5E** and **5F**). Although the MULTICOM series remained among the top performers, Kiharalab dropped in the rankings, while Zheng and Zheng-Multimer rose into the top tier. Kiharalab’s decline appears to stem primarily from its failure to select strong first models in Phase 0, likely due to incorrect stoichiometry predictions. We also ranked groups using Phase 0 and Phase 2 targets, respectively, using both best models and first models (supplemental **Fig. S8** and **S9**). These results show that top-performing groups are mostly robust in their performance and consistently rank near the top across phases. However, in each single phase, the statistical significance for performance differences among different groups is weak.

The majority of the top 25% groups, with the exceptions of kozakovvajda and CoDock, relied on AF2/3-based frameworks as their core modeling engine. Top groups such as the MULTICOM series, Kiharalab, and Yang-Multimer all used AF3 in combination with large-scale sampling. At present, the challenge appears to be shifting from method innovation toward a competition of scale—where performance is increasingly affected by sampling effort, raising concerns about the diminishing role of conceptual methodology innovation.

### Insights from additional CASP16 experiments

Beyond evaluating group performance, a central goal of CASP is to understand the progress achieved and the challenges that remain. To identify key factors influencing prediction accuracy, we examined performance across different phases.

In Phase 0, predicting the correct stoichiometry becomes a primary challenge. Many groups chose to sample different stoichiometries with their five models. Because there are a limited number of possible stoichiometries for most targets, it is easier to obtain the correct one among five predictions. Thus, we only focused on the first models and considered the stoichiometry in model 1 as the prediction for each group. **Fig. 6A** shows stoichiometry prediction accuracy per target for each group. Top-performing groups correctly predicted stoichiometry for most targets (above 70%). However, it’s important to note that most targets involved simple stoichiometries such as A1B1, which can be easily predicted based on the dominance of such stoichiometry among experimentally determined protein complexes. Targets with higher and rarely observed stoichiometries, like H1265, were poorly predicted.

**Figure 6.**
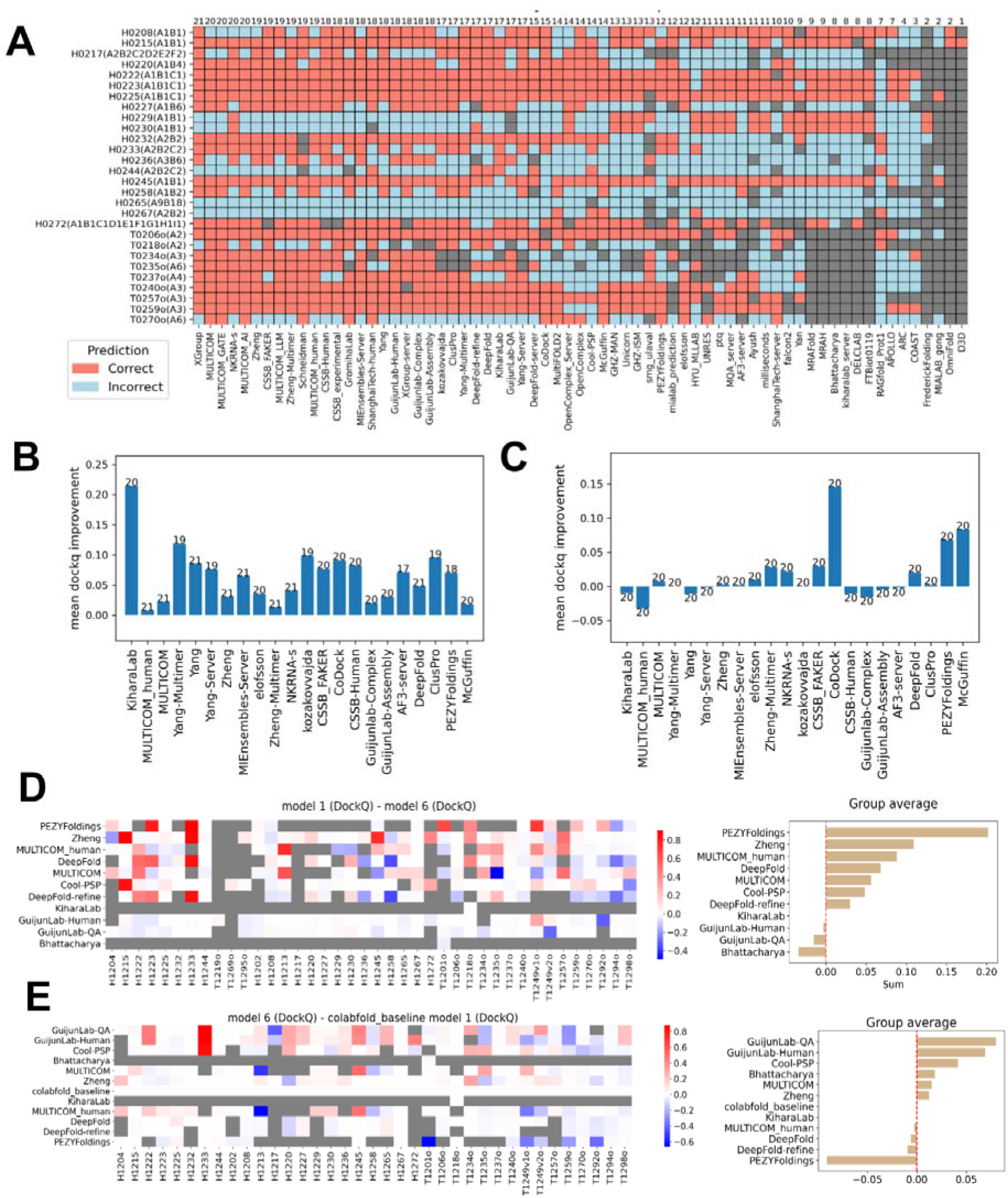
Performance differences across phases and model 6 evaluation. **A)** Stoichiometry prediction accuracy for each group across Phase 0 targets, red: correct, cyan: incorrect, grey: no value. **B)** Average DockQ improvement for each group from Phase 0 to Phase 1. **C)** Average DockQ improvement for each group from Phase 1 to Phase 2. In both **B** and **C**, we used the RBM pipeline to compute DockQ scores, making scores comparable between phases. **D)** Left: per-target DockQ difference between the first submitted model and model 6 (where model 6 was generated using ColabFold MSAs). Right: sum of DockQ differences for each group. **E)** Left: per-target DockQ difference between model 6 and the first model from Colabfold. Right: Sum of DockQ differences for each group.

Interestingly, two targets with simple stoichiometry, H0229 and H0230, were mispredicted by many groups. H0229 and H0230 are homologous and they share a common PDB template, 7rw6, which shows high sequence identities to H0229 (40%) and H0230 (80%). Notably, 7rw6 has a stoichiometry of A2B2, matching many submitted predictions. This suggests that some groups may have inferred stoichiometry by homology to complexes in the PDB. For H0267, another target for which only two groups predicted stoichiometry correctly, several templates, such as 8it0 and 8ffi, are available. However, these templates differ in stoichiometry, preventing homology-based stoichiometry inference. It is possible to predict stoichiometry *de novo* by trying to model a protein complex at varying stoichiometry. For example, target T0235o, a hexamer with no detectable templates via HHblits [37], was correctly predicted by many groups.

We evaluated the first models of different groups in Phase 0 and Phase 1 using the RBM pipeline, which will penalize incorrect stoichiometry predictions that lead to incorrect PPI interfaces. As expected, we observed a robust improvement in prediction accuracy of model 1 from Phase 0 to Phase 1 (**Fig. 6B**). Over half of the top 22 (25%) groups showed average improvements in DockQ scores of greater than 0.05, suggesting that knowing stoichiometry substantially benefited interface prediction accuracy. In contrast, the transition from Phase 1 to Phase 2, where MF models were provided, stimulated limited performance improvement (**Fig. 6C**). This is likely related to the fact that many of the top groups did massive sampling using their own pipelines. Additionally, MF mostly uses AFM to model, while the CASP16 groups can use the AF3 server to generate models of slightly higher quality. Only three groups, CoDock, PEZYFoldings, and McGuffin, achieved mean DockQ improvements greater than 0.05 from Phase 1 to Phase 2. PEZYFoldings also showed the best performance in selecting the first model among their predictions, and thus, they might have a higher ability to select better predictions from MF models.

CASP16 also introduced a new experiment, so-called “model 6”, aimed at disentangling the contributions of MSA quality and protein modeling deep learning networks. Participants were asked to generate models using ColabFold MSAs, thereby standardizing the input MSAs. We reason that the comparison of model 6 against each group’s model 1, assuming the same modeling pipeline was used for both, reflects their ability to obtain better input MSAs than ColabFold. In contrast, the comparison of model 6 against ColabFold’s model 1, because the same input MSA was used, reflects the performance of a group’s modeling pipeline independent of input MSAs. Although participation in this challenge was limited—only 11 groups submitted results, and two of them only for one or two targets—the comparative analysis still yields preliminary insights and considerations for the design of future CASP challenges.

For most groups that participated in the “model 6” challenge, their first models outperform their “model 6s”, suggesting that many groups customized or refined their MSAs to improve prediction quality (**Fig. 6D**). However, an alternative explanation is that participants invested more effort into sampling and model selection for their first model, while model 6 may have been produced with minimal sampling and less rigorous ranking. Comparing each group’s model 6 against ColabFold’s model 1 revealed that most groups showed performance improvement over ColabFold (**Fig. 6E**). This is not unexpected, but it is hard to judge how much such improvement is simply achieved by broader model sampling, a simple strategy known to improve the chance of obtaining a better prediction.

One notable group from both analyses was PEZYFoldings. Although they showed the largest improvement from model 6 to model 1, their model 6 is worse than ColabFold’s model 1. According to their methodology descriptions from CASP15, PEZYFoldings enhanced their MSAs using short-read archive data to increase sequence depth and applied PLMs to improve alignment quality. For modeling, they used AF2 or AFM and generated 1,000–2,000 models per target. In contrast, for model 6, they only applied PLM-enhanced MSAs, generated five models using AFM, disabled template usage, and selected the best among them. It is plausible that disabling templates slightly reduced the quality of their model 6, while the improved MSA and large-scale sampling contributed to the substantial gain observed in their model 1 over model 6. This observation also supports our earlier hypothesis that groups may have invested less effort in model 6 compared to their primary submissions. Unfortunately, most other groups did not provide sufficient detail on how their model 6 was generated. As a result, it is difficult to disentangle the specific effects of MSA quality, sampling depth, and model ranking. This lack of technical details limits our ability to draw conclusions from the above analyses.

### Model ranking and selection

Having proven to be a winning strategy in CASP15, massive model sampling was adopted by many top-ranking groups in CASP16. Methods like MF made it relatively easy to generate a diverse set of models, and the diffusion module used in AF3 also made the system well-suited for model sampling. Therefore, we expect that methods for model ranking and selection will become an important focus for future development. To directly evaluate each group’s ability in model ranking, we calculated the percentage of targets for which a group’s first model was also the best among the five submitted models. Among the top-ranking groups, many performed better than random selection, where the first model would have a 20% chance of being the best (**Fig. 7A** and **7B**). However, model ranking and selection remain major challenges for the field. Most top-ranking groups selected the best model first for less than 40% of targets, and AF3 selected the best model first in only 25% of cases. The one notable exception was the PEZYFoldings group, which succeeded in selecting their best models as first models for 58% of targets.

**Figure 7.**
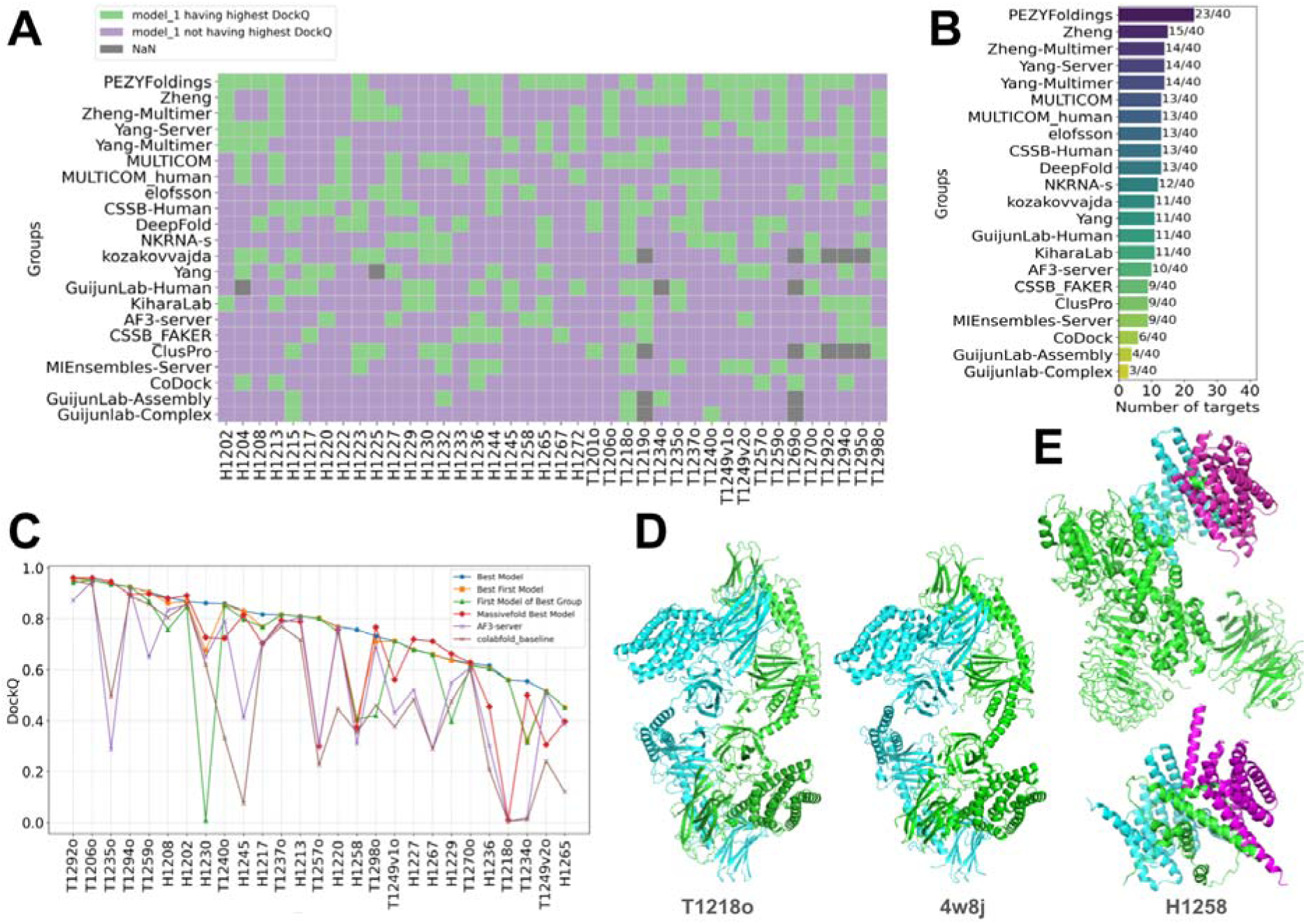
Evaluation of model ranking ability and the community’s improvement over MF. **A)** Evaluation of whether the first model selected by a group is the best model for Phase 1 targets and top-performing groups (in overall ranking over all phases). Green: the first model being the best model; Purple: the first model is not the best model. **B)** The number of correctly picked first models (when the first model is the best for a target) by each group shown in **A**. **C)** Values of the highest DockQ among MF models or different sets of models submitted by CASP16 groups. Targets are ordered by the highest DockQ among models submitted by CASP16 groups from left to right. **D)** An example, T1218o, where CASP16 groups using docking-based methods outperformed AF2/3-based predictions. AF-based predictions were poor, although a homologous template (right) is available for this target. **E)** An example, H1258, where the Yang lab can achieve significantly higher interface quality by only including the segment of a protein (the green one) that mediates its interaction with others in the model. Top: the target; bottom: a model from the Yang lab.

PEZYFoldings demonstrated superior performance in model ranking for both monomer [cite: our monomer paper] and oligomer targets, and their strategy may serve as an inspiration for other groups. According to their method description, the PEZYFoldings group coupled model refinement with model ranking, and we encourage researchers in the field to study their manuscript [38] and learn from their experience.

Utilizing the large number of predicted structures generated by MF for many targets, we investigated the following question: Would it be possible to obtain better predictions among the MF models if a perfect model selector were available? To address this, we ran OST for all MF models against their corresponding targets. We then plotted the best DockQ score from the MF models for each target against the best models submitted by CASP16 groups for the same targets (**Fig. 7C**). AA targets, which were separately analyzed in a previous section, were excluded. Overall, the best models generated by CASP16 groups were generally better than the best MF models. On many targets, including H1230, T1240o, H1217, T1257o, H1258, T1249v1o, T1249v2o, H1236, and T1218o, the CASP16 groups’ best models substantially outperformed the best MF models. Only for a few targets did the pool of structures generated by MF contain better models.

This observation explains why the performance improvement from Phase 1 to Phase 2 was not obvious for the top-performing groups. The pool of models sampled by CASP16 groups was of higher quality than those generated by MF. Notably, MF predictions were produced prior to the public release of AF3 and were generated using various sampling methods based on AFM. In contrast, top-performing CASP16 groups were able to sample a larger number of models using both AFM and AF3, and the additional MF models did not significantly impact their performance. As a result, the significance of the CASP16 Phase 2 experiment was diminished, highlighting the need for a better design of this phase in future CASP experiments. For example, rather than asking groups to submit their own models in Phase 2, it may be more meaningful to ask participants to select the best five from a large set of models for each target which, in fact, was one of the quality assessment (QA) experiments in CASP16. However, we observed that not all participants in the protein oligomer prediction challenge took part in the QA experiment. Notably PEZYFoldings, which appeared to perform best in model selection, did not participate in QA experiments. Thus, a strategy to encourage as many as participants to join the QA is needed.

Nevertheless, this analysis allowed us to identify interesting targets where the field’s best predictions substantially outperformed MF’s best models in interface prediction accuracy. One notable case is target T1218o, a dimer of the delta-endotoxin from *Bacillus thuringiensis*. For this target, the community outperformed both MF and AF3, with the best submitted model reaching a DockQ score of 0.56 (AF-generated predictions have DockQ close to 0). A homologous template (PDB: 4w8j) was available for T1218o, showing decent sequence similarity (36% identity, 92% coverage) and high oligomeric interface similarity (**Fig. 7D**). However, this template was likely not properly utilized by AF2 or AF3. Structural analysis suggests that the biologically relevant dimeric state of 4w8j can only be obtained by generating symmetry mates and selecting a dimer with a large interface area—an assembly not included in the biological assembly information provided in the PDB (mmCIF format). As a result, template search procedures in AF2/AF3, which likely did not consider symmetry-derived oligomers, failed to identify an appropriate template, leading to poor performance for groups relying on AF-based modeling (DockQ scores close to zero for AF3-server). Interestingly, the top three models for T1218o were submitted by groups—FTBiot0119 [cite: their paper if available], kozakovvajda, and ClusPro—that employed docking-based approaches rather than AF2-based modeling. The only non-docking group showing comparable performance (DockQ > 0.5) was DeepFold-server [cite: their CASP paper], which implemented a careful pipeline focused on template selection and refinement of template–target alignments.

Another notable example is H1258, a human LRRK2/14-3-3 complex mentioned above. The best models submitted by Yang-Multimer and Yang-Server, focused only on residues 861–1014 of LRRK2 and two copies of 14-3-3, exactly where the interaction happens, achieving DockQ scores above 0.75 (**Fig. 7E**). In contrast, when they modeled the full-length LRRK2, interface accuracy dropped to about 0.4, on par with other groups and MF predictions. This case illustrates how one can carefully refine the sequences used for modeling and improve interface prediction. Constraining to the regions mediating the interactions possibly helps to reduce the complexity AFM/AF3 needs to handle, allowing them to focus on getting a better interface. Refining the boundary of a construct is a commonly used strategy in experimental structural biology, and its application in computational structural biology should be further explored.

### Evaluation of hybrid targets

Following the major accuracy improvements in protein assembly prediction observed in CASP15 and the expansion of methods to predict biomolecular interactions involving nucleotides and small molecules [14,39,40], CASP organizers have been actively developing new challenges to drive further innovations in the field. As part of this effort, the hybrid target category was expanded from 2 targets in CASP15 [41] to 16 in CASP16. These targets consist of both proteins and nucleotides (DNA and RNA). Compared to other types of non-protein molecules found in experimental structures, DNA and RNA ligands are relatively large in volume, making it crucial to model the protein and DNA/RNA components of these complexes jointly.

There were 16 hybrid targets in Phase 1, with targets M1228 and M1239 each having two possible conformations. Three of these hybrid targets were also used in Phase 0. However, since this was the first time hybrid targets were included, participation was limited. CASP16 received 2,226 models from 57 groups for Phase 1 and 392 models from 33 groups for Phase 0. Many groups submitted models for only a small number of hybrid targets, with only 23 groups in Phase 1 submitting models for at least 9 targets. Interestingly, for hybrid target M1268, more groups participated in Phase 0 than in Phase 1. The correct stoichiometry of M1268 is A8B8R8V8W1X1; in Phase 0, every group predicted a much smaller complex without knowing this information. Modeling the complexes with high molecular weight using the correct stoichiometry might have been challenging for many teams due to limitations in GPU memory.

We adapted the OST and RBM pipelines described above to evaluate the Phase 1 and Phase 0 hybrid targets, respectively. Three measures, QSbest, DockQ and TM-score, were omitted due to the following reasons. First, the QSbest was not designed for nucleotides and not fully adapted to evaluate hybrid targets according to the OST developer. Second, the current OST pipeline used by the Prediction Center failed to generate DockQ scores for hybrid targets, preventing the inclusion of DockQ in the OST pipeline. Third, although US-align could compute TM-scores for protein-NA complexes, we found it might fail to align NA chains even if we used it to align the target structure to itself. The remaining three scores (ICS, IPS, and lDDT) were used, and z-scores of them were weighted equally to generate the cumulative z-scores. We also separated the protein-protein interfaces and protein-NA interfaces, and used the two interface scores (ICS and IPS) to evaluate their quality. NA-NA interfaces are evaluated elsewhere in this issue [cite casp16 RNA assessment].

**Fig. 8A – 8F** summarizes group performance across three categories: full models **(Fig. 8A** and **8B**), protein–protein interfaces (**Fig. 8C** and **8D**), and protein–nucleic acid interfaces (**Fig. 8E** and **8F**). Similar to the protein assembly challenge, no clear winner showing significantly better performance than other groups emerged in this category (**Fig. 8B, 8D,** and **8F**). Kiharalab ranked first in the full model category, MIEnsembles-Server from the Zheng team ranked highest for protein–protein interfaces. For protein-NA interfaces, three groups, Vfold, Kiharalab, and MIEnsembles-Server, ranked highest with very similar cumulative z-scores. Notably, the AF3 server ranked well behind the top-performing groups. However, the superior performance of these groups over AF3 is likely attributed to more extensive modeling and selection from larger pools of models. Several of the top-ranking groups, including Kiharalab, Vfold [cite: Vfold paper], and CSSB_experiment, utilized AF3 for hybrid targets, while three groups from the Zheng lab, MIEnsembles-Server, NKRNA-s, and Zheng, did not rely on AF3 but instead used deep learning networks derived from AFM, demonstrating that competitive performance in modeling hybrid complexes can be achieved through adapted AFM-based approaches.

**Figure 8.**
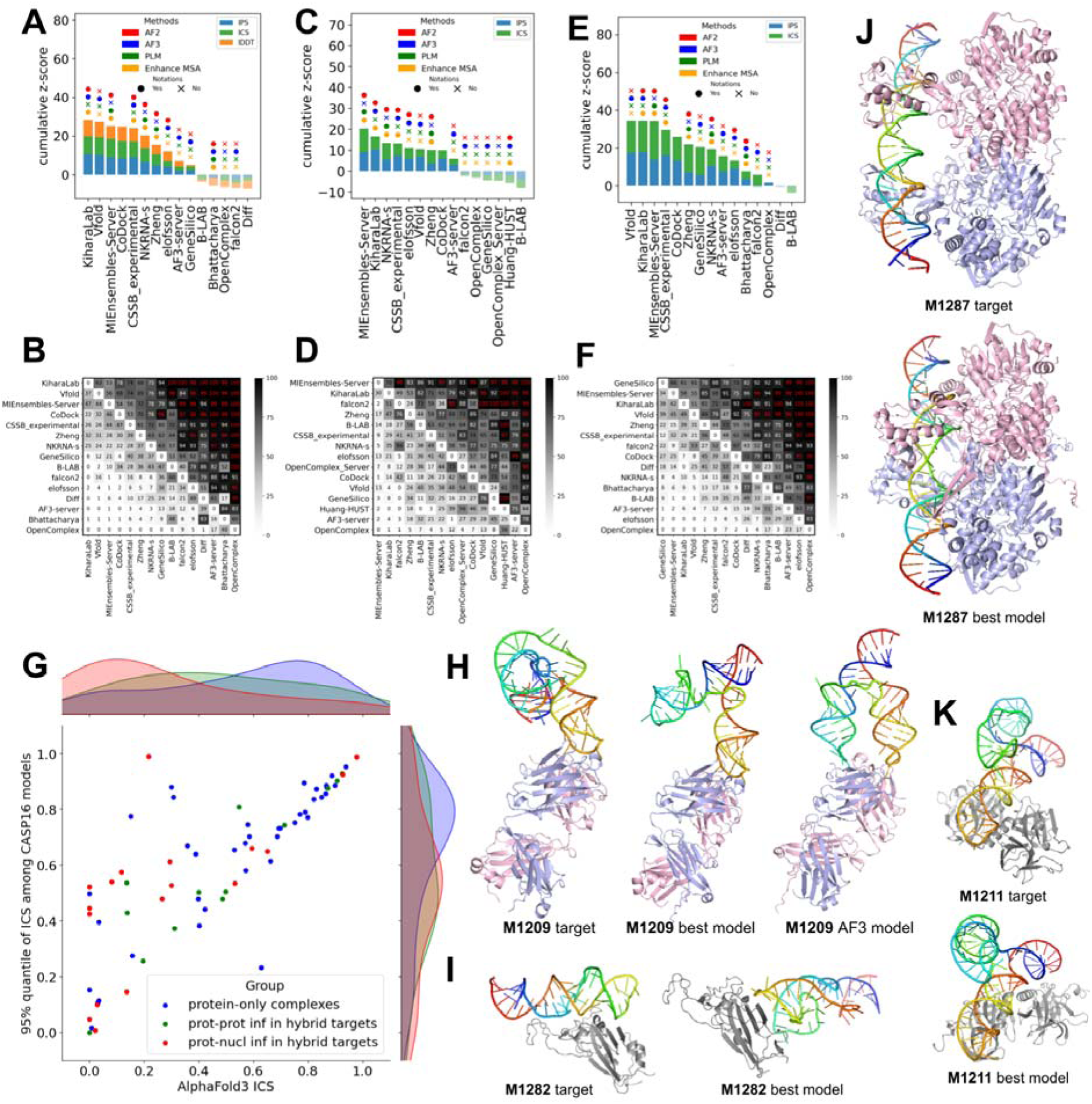
Evaluation of hybrid targets. **A-F)** Ranking and head-to-head bootstrap comparisons of prediction accuracy for full models (**A** and **B**), protein-protein interfaces (**C** and **D**), and protein-NA interfaces (**E** and **F**), respectively. **G)** Comparison of AF3 ICS scores with the 95th percentile ICS scores among CASP16 submissions for different target interface types. Blue: protein-only complexes; green: protein-protein interfaces in hybrid targets; red: protein-NA interfaces in hybrid targets. Density plots along the axes show the distribution of ICS scores for each interface group. **H)** Structures of target M1209 along with the best model from participants and the best AF3_server model. **I-K)** Targets (M1282, M1287, and M1211) and their corresponding best models. These are difficult targets lacking predictions of acceptable quality (best ICS for protein-NA interface < 0.2).

Compared to protein-only assemblies, hybrid targets remain more challenging. Protein–protein interfaces achieved higher prediction accuracy (as measured by ICS) in protein-only targets for both AF3-server and participants compared to protein–protein interfaces within hybrid targets (**Fig. 8G**, green vs. blue). In contrast, protein–NA interfaces were more difficult for the AF3-server to predict than protein–protein interfaces in hybrid targets, whereas top-performing groups achieved comparable performance for both protein–NA and protein–protein interfaces (**Fig. 8G**, red vs. green). However, because participants employed extensive model sampling while the AF3 server produced only five models per target, direct performance comparisons between AF3 and other groups should be interpreted with caution.

**Fig. 8H** presents an example, M1209, where top-performing groups produced better models than AF3. In this target, the fragment antigen-binding region (Fab) which, for crystallization, replaces the IIc loop from the HIV-1 RRE stem loop II [42]. This Fab-RNA loop has previously been used to crystalize other RNA molecules, so as expected, participants accurately predicted a small, specific interface between the RNA and the antibodies. Whereas the AF3 model incorrectly bent the RNA to form an enlarged interface. Targets (M1282, M1287, and M1211) for which the community failed to predict accurate models, along with the best models by ICS, are shown in **Fig. 8I – 8K**. For these targets, not only were the protein-NA interfaces incorrectly predicted, but the conformations of the NAs were also poorly modeled. While single-domain protein structure prediction is largely considered a solved problem, modeling 3D structures of NAs remains a major challenge for the field [cite: Rhiju’s paper]. Similarly, modeling protein-NA hybrid complexes is also more challenging than modeling protein-only complexes.

## Conclusions and Future Perspectives

CASP16 highlighted both steady progress and persistent challenges in the field of protein complex structure prediction. Contrary to monomer structure prediction, oligomer structure prediction remains an unsolved problem. Overall, the performance of participating groups improved moderately relative to CASP15. Notably, top-performing groups such as Kiharalab, the MULTICOM series, and Yang-Multimer consistently produced high-quality models across multiple evaluation metrics, demonstrating the effectiveness of combining AFM or AF3 with customized MSAs, extensive model sampling, and modeling construct refinement. By iteratively refining the sequence range included in modeling constructs, the Yang lab was able to generate higher-quality interfaces for several targets that were particularly challenging for other groups. We expect that this strategy will drive further improvements in future CASP experiments.

AF3, released at the beginning of CASP16, was a major contributor to the observed progress in the experiment. Although the AF3-server with standard parameters appeared to fall behind many CASP16 groups, most top-performing groups incorporated AF3 into their pipelines. Extensive model sampling and selection from large pools of models enabled these groups to outperform the AF3-server, which submitted only five models per target. Now that the AF3 source code has been released, we expect researchers in the field will further optimize its performance by refining MSAs and input constructs.

Beyond AF3, we observed other promising approaches that could drive future progress in the field. One such approach is the use of traditional protein–protein docking methods for AA complex prediction. The kozakovvajda group emerged as a dominant performer in this category, delivering accurate predictions for 60% of targets that were considered challenging for AF2/3 and other leading methods in CASP15. These results demonstrate that for particularly difficult targets—those lacking evolutionary signals and involving small interface regions—carefully designed docking strategies and sophisticated sampling pipelines can lead to substantial gains in accuracy. Due to the limited number of AA targets, it remains difficult to draw definitive conclusions about overall progress in this category since the last CASP. However, as AF-based methods continue to dominate the field, the outstanding performance of the kozakovvajda group highlights the promise of alternative, physics-based approaches and may encourage the community to further explore modeling strategies that diverge from the AF paradigm.

Another promising development is the model ranking and selection pipeline used by the PEZYFoldings group, although they unfortunately did not participate in the quality assessment challenge. As massive model sampling becomes a dominant strategy, the ability to accurately rank predictions is becoming increasingly crucial. To stimulate progress in this area, CASP16 introduced Phase 2, which provided a large number of MF models to help predictors obtain better models. Model ranking remains a weak point for most groups, and even the top-performing groups selected their best models as first models for only 30%–40% of targets. However, by employing a pipeline that couples model ranking with model refinement, the PEZYFoldings group performed markedly better than others in selecting first models in both the monomer and oligomer modeling categories.

Due to the relatively small number of participants, the limited number of targets, and the fact that established quality evaluation software does not function robustly on protein-NA hybrid targets, less emphasis was placed on evaluating these targets in CASP16. However, this emerging category represents an important unsolved challenge for the community. Predicting the conformations of nucleic acids and their interfaces with proteins remains difficult even for the state-of-the-art methods. Although the current top-performing groups in this category relied on AFM/AF3, we expect this area to offer significant opportunities for innovation and progress in the near future.

Looking ahead, several methodological challenges remain. First, there is an urgent need for improved scoring functions to enhance model selection. The pool of sampled structures submitted by CASP16 groups collectively contained correct models for the majority of oligomer targets, yet no single group was able to consistently select the correct model across targets. Second, accurate prediction of AA complexes remains difficult, although CASP16 demonstrated the potential of docking-based approaches in such cases. Third, while AF-derived methods have dominated recent CASP experiments, template-based modeling and docking strategies continue to provide value for difficult targets (such as T1218o) and should not be overlooked. Fourth, the Phase 0 experiment of CASP16 shows that it is possible to predict stoichiometry based on homology and comparative modeling across different stoichiometries; however, predicting high-order and complicated stoichiometries remains challenging. Finally, we hope that greater emphasis will be placed on developing conceptually innovative methods rather than relying primarily on incremental improvements over AF.

### Notes for future CASPs

Because oligomer structure prediction has shown steady progress in recent CASP experiments, we expect it to remain an important category in the next CASP. Some lessons learned from evaluating CASP16 may be useful for future organizers and assessors. **First**, the Phase 0 experiment, which tests the ability to predict structures solely from sequences, is particularly valuable, and we hope that this challenge will be emphasized in the next CASP. **Second**, fairly evaluating Phase 0 targets, where the target and the models may have different stoichiometries, can be challenging. We introduced the RBM pipeline to address this issue, and we hope that our approach will inspire future assessment strategies. **Third**, care should be taken in interpretation results for AA target AA targets. When multiple antibody or antigen chains are present, OST-based scores may be dominated by antibody–antibody or antigen–antigen interfaces, rather than antibody–antigen interfaces. To better emphasize the intended AA interactions, special attention needs to be paid to such cases. **Fourth**, evaluation methods for protein–NA hybrid targets are still poorly developed. For example, DockQ and QSbest were not fully adapted to evaluate hybrid targets, and USalign [43] is also not robust for these targets. To establish more reliable evaluation methods, it would be beneficial to recruit a dedicated assessor for this category in the next CASP. **Fifth**, the Phase 2 experiment was poorly designed, as providing MF models was not helpful for many groups that performed large-scale model sampling independently, especially using AF3. To better stimulate progress in model selection, we suggest revising Phase 2 so that groups select five models from a large pool provided by the organizers, with evaluation merged into the quality assessment category. **Finally**, to some extent, the major challenges in CASP16 have shifted toward a competition of computational scale, where extensive model sampling using AF2/3 has become the dominant winning strategy. This trend risks discouraging conceptual methodological innovation. We believe future CASP experiments should refocus on areas where AF-like networks may not represent the optimal approach, such as AA target modeling, model ranking, stoichiometry prediction, and protein–NA target modeling.

## Methods

### Refining target definitions

Originally, CASP16 provided participants with over 40 oligomer targets, including the alternative conformations for multiple complexes. However, upon closer examination, we removed some targets representing alternative conformations. T1294v1o and T1294v2o were supposed to represent two distinct states. However, because the structures of these targets are nearly identical (RMSD < 0.5), only T1294v1o was kept. In contrast, for T1249, both versions were retained and treated as separate targets in the assessment. This decision was based on the target description, which explicitly informed participants of the presence of distinct open (v1) and closed (v2) states and instructed them to submit models for each state. Substantial structural differences between the two states also justified their separate inclusion in the assessment.

For target T1295o, the original target description defined it as an 8-mer (A8). However, the biological assembly revealed in the experimental structure has a stoichiometry of A24, and each chain is a fusion between an antibody and a fimbrial adhesin—the structures of both were previously known. However, the default OST evaluation pipeline produced low scores for every model, mostly because the relative orientation between the antibody and the adhesion was poorly predicted. Manual inspection indicates that the relative orientation between these two fused segments should be flexible, and the solved structure is likely an average of possible conformations. Therefore, we trimmed the antibody portions and defined the 24-mer adhesin as the target. Although this structure had already been solved and deposited in the PDB, we retained it in the challenge. Due to the difference in stoichiometry between the target (A24) and the Phase 1 models (A8), we used our RBM pipeline for its evaluation.

### The overview of targets, groups, and factors affecting prediction

To visualize target properties, group performance, and key methodological features, we generated a heatmap (**Fig. 1A**) of the best DockQ score of each group for each target, excluding groups that submitted models for fewer than 25% of targets. To reveal performance patterns, Ward’s hierarchical clustering algorithm was applied to both predictors and targets based on DockQ scores. Annotations describing predictor strategies and target characteristics were obtained from participant abstracts and official CASP target descriptions, respectively.

To investigate factors influencing prediction performance, we computed the Neff for each interface based on the *colabfold_baseline* MSAs. Interfaces within each target were partitioned into homo-dimeric and hetero-dimeric categories based on chain mapping information provided by the OST pipeline. For each homo-dimeric interface, we extracted positions corresponding to the single protein from the MSA; for each hetero-dimeric interface, we extracted positions corresponding to the protein pair from the MSA, retaining only sequences that contained non-gap positions for both proteins. Additionally, we defined complex size as the total number of residues in the assembly, while interface size was measured by the number of interacting residue pairs (residues from different molecules within 5 L of each other).

### Evaluations and ranking

For a comprehensive evaluation, we included **ICS**, **IPS**, **DockQ**, and **QSbest** to assess interface accuracy, and **TM-score** and **lDDT** to evaluate global fold and local geometry. The inclusion of TM-score and lDDT is one of features that distinguishes CASP from CAPRI, which mainly focuses on interface evaluation. A description of each score is shown in **Table 1**. As explained above, we only used ICS, IPS, DockQ, and QSbest for AA targets to focus on antibody–antigen interface quality; due to technical difficulties of applying certain software to protein-NA complexes, we only used ICS, IPS, QSbest, and TM-score for hybrid targets; we used all six metrics for the remaining “normal” targets.

**Table 1.** Scores used for oligomer target evaluation.

| Score name | Evaluation aspects | Normal | AA | Hybrid | Description |
| --- | --- | --- | --- | --- | --- |
| ICS | interface accuracy | yes | yes | yes | ICS is an F1-score representing the harmonic mean of the precision and recall of interacting pairs.<br>ICS = $F1 = 2 * \text{precision} * \text{recall} / (\text{precision} + \text{recall})$ |
| IPS | interface accuracy | yes | yes | yes | IPS is the Jaccard coefficient of interface residues between models and targets. $IPS = (M11) / (M10 + M10 + M11)$<br>**M01**: the number of the model interface residues that are not present in the target interface<br>**M10**: the number of the target interface residues that are not present in the model interface<br>**M11**: the number of interface residues that are present both in the model and the target interface |
| DockQ | interface accuracy | yes | yes | no | The composite score of $F_{\text{nat}}$ (how many native inter-chain contacts are preserved), LRMS (deviation of the ligand after superimposing the receptor), and iRMS (root mean square deviation over interface residues) |
| QSbest | interface accuracy | yes | yes | no | QS-score measures the similarity of shared interface contacts, normalized by the total number of interface contacts in model and target. |
| TM-score | global fold | yes | no | no | Similarity between the model and the target based on their optimal superposition |
| IDDT | local geometry | yes | no | yes | IDDT is the average fraction of local interatomic distances in the target that are preserved in the model within thresholds of 0.5, 1.0, 2.0, and 4.0 Å. |

As described in the Results section, for cases where the models shared the same stoichiometry as the target—i.e., most Phase 1 and Phase 2 targets—we used the outputs of OpenStructure (OST, v2.9.2) provided by the CASP organizers to extract evaluation scores. We directly took the overall TM-score and lDDT values from the OST outputs. For interface quality scores (ICS, IPS, DockQ, and QSbest), instead of using the overall scores computed by OST across all interfaces within a target, we extracted the score for each individual interface and developed a weighted scoring scheme to aggregate scores across the entire assembly (see equation below). Specifically, by using the base-10 logarithm of the interface size as the weight, our formula prevents large interfaces from disproportionately dominating the final scores. Larger interfaces tend to be easier to predict, while smaller interfaces provide better discrimination among CASP groups. At the same time, since smaller interfaces have a higher chance of being experimental artifacts and are generally less critical for complex formation, we weighted them less heavily rather than assigning equal weight to all interfaces.

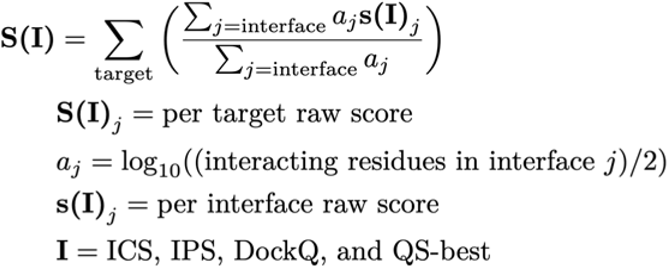

For Phase 0 targets, as well as T1269o (fiber), T1219o (fiber), and T1295o, where the targets differ from their models in stoichiometry, we used the RBM pipeline described in the Results section. RBM computes the best ICS, IPS, QSbest, DockQ, TM-score, and lDDT values for each interface in the target and model. Interfaces were identified based on OST outputs, and for each interface in the target, we identified its best matching interface in the model by maximizing each score. The same procedure was repeated for each interface in the model. ICS, IPS, and QSbest were computed using in-house scripts based on their definitions described in Table 1, with contacts between chains defined as residue pairs within 5 L. DockQ, lDDT, and TM-score were computed using established scripts [8,18,43]. Per-interface scores were weighted by the base-10 logarithm of the interface size, following the weighting scheme described above. The weighted averages of per-interface scores were computed separately for the target and the model, and the final score for each target–model pair was obtained by averaging these two weighted scores.

Following the tradition of CASP, we used Z-scores of each metric for performance evaluation and ranking. We weighted the z-score for each metric equally to obtain an average z-score for each model. We noticed cases in which certain models could not be evaluated for specific metrics due to intrinsic model problems. In such cases, missing values were imputed with the lowest observed value across all models for that target and metric. Z-score calculations were conducted in two rounds. In the first round, models with an initial Z-score below −2 were considered outliers and excluded. Z-scores were then recalculated, and in the final rankings, any Z-scores below −2 were imputed using −2. This approach prevents groups using more experimental or high-risk strategies—who may occasionally generate poor models—from being excessively penalized and allows consistent high performance on other targets to be recognized.

As in previous CASPs, final rankings were determined using cumulative z-scores across targets. Phase-specific rankings were calculated by summing ZLscores across targets within each phase, while overall rankings were calculated by combining ZLscores across all targets from the three phases. Rankings were generated for both **first models** and **best models** (the model with the highest average z-score over all metrics) for each group. If a group did not explicitly designate its first model, the model with the lowest numerical model ID was used.

To assess the robustness of the rankings and evaluate whether observed performance differences between groups were statistically significant, we performed head-to-head comparisons using only the common targets for which both groups submitted predictions. We generated 1,000 bootstrap replicates of these common targets. For each replicate, pairwise comparisons were performed based on the cumulative z-score over the bootstrap sample, and the number of replicates in which each group outperformed the other was recorded to estimate statistical significance.

### Progress evaluation

To enable comparison with CASP15, we required a baseline predictor with consistent performance across both CASP rounds. ColabFold, which participated in CASP15, served as such a reference. Its developers—Dongwook Kim, Seongeun Kim, and Milot Mirdita— generously provided models generated using the similar CASP15 pipeline with updated databases and AFM weights (from multimer-v2 to multimer-v3), enabling a consistent baseline for assessing community-wide progress. In addition, the AF3 server became publicly available shortly after the release of CASP16 targets. Many participants indirectly incorporated AF3 into their prediction pipelines. Thus, AF3 also serves as a reference to evaluate whether participants were able to surpass its default performance. Thanks to the Elofsson group, all CASP16 targets were submitted to the AF3 server, and the resulting models were submitted as the AF3-server group (group number: TS304).

The same analysis pipeline was applied to both CASP15 and CASP16 results. For each target, the model with the highest overall DockQ (not the per-interface values, to be consistent with CASP15) from each group was selected. Groups that submitted fewer than 25% of the targets were excluded. Targets missing from the *colabfold_baseline* were also removed. To fairly compare performance between CASP15 and CASP16. We used two statistical methods to select a set of CASP15 targets showing a similar difficulty level distribution to CASP16 targets. We estimated the difficulty of a target by the DockQ score of *colabfold_baseline’*s first model, and we used a Hungarian algorithm to generate a one-to-one mapping between CASP15 and CASP16 targets, while minimizing the difficulty difference between members in the two sets. To introduce randomness into this process, we first used Gaussian Kernel Density Estimation to fit a continuous probability distribution for CASP15 and CASP16 target difficulties (by DockQ of *colabfold_baseline* models). Then, a weighted resampling based on the ratio between the two distributions is done to ensure the bootstrap sample from CASP15 resembles CASP16 in target difficulties.

For AA complexes, we followed the same treatment as in CASP15: antibody chains were manually concatenated into a single chain, as were antigen chains, to form one unified antibody–antigen interface. This approach ensures comparability between CASP15 and CASP16. All DockQ scores were computed using the same OST version (v2.9.2). Except for the CASP15 comparison shown in **Fig. 4A – 4F**, DockQ values used in other analyses represent the average over all antibody–antigen interfaces without chain concatenation. Because AF3’s improved performance on AA targets depends on massive sampling using different random seeds, we performed extensive AF3 modeling of the AA targets using a standalone version of AF3 downloaded from the GitHub repository of DeepMind. For each AA target, we generated 5,000 models using 1,000 random seeds. We ranked these models by average ipTM of the antibody–antigen interfaces in a target, and the model with the highest ipTM was considered the best prediction. We used the same CASP15-like pipeline to evaluate the quality of these best predictions. To evaluate the performance of the standalone AF3 with single seeds, we also used the first models generated by each seed and averaged their DockQ scores.

## Author Contributions

JZhang and RY conducted the assessment with supervision from QC, and with instructions from AK and GS. RDS, JZhou, NVG, RCK, and RD participated in data analysis and provided suggestions for analysis methods. JZhang, RY, and QC prepared the figures and drafted the manuscript. All authors edited the manuscript.

## Acknowledgements

QC is supported by the Endowed Scholars Program in UTSW and I-2095-20220331 from Welch Foundation. NVG is supported by I-1505 from Welch Foundation and 2224128 from National Science Foundation Division of Biological Infrastructure. RDS is supported by NIGMS GM147367, JZhou is supported by RR190071 from the CPRIT and 4DP2GM146336-02 from NIGMS, and JZhang is supported by NIAID K99AI180984-01A1. AK is supported by NIGMS GM100482. The authors thank all CASP organizers, particularly John Moult, for their valuable guidance during the CASP evaluation. We are also grateful to Gabriel Studer for instructions on using OST, to CAPRI assessor Mark Lensink for discussions on target partitioning and for sharing CAPRI results, to Guillaume Brysbaert for providing the MassiveFold structures, and to Arne Elofsson for his feedback and for identifying several missing models during our evaluation.

