## Supplementary material for "Assessment of Protein Complex Predictions in CASP16: Are we making progress?": Fig. S

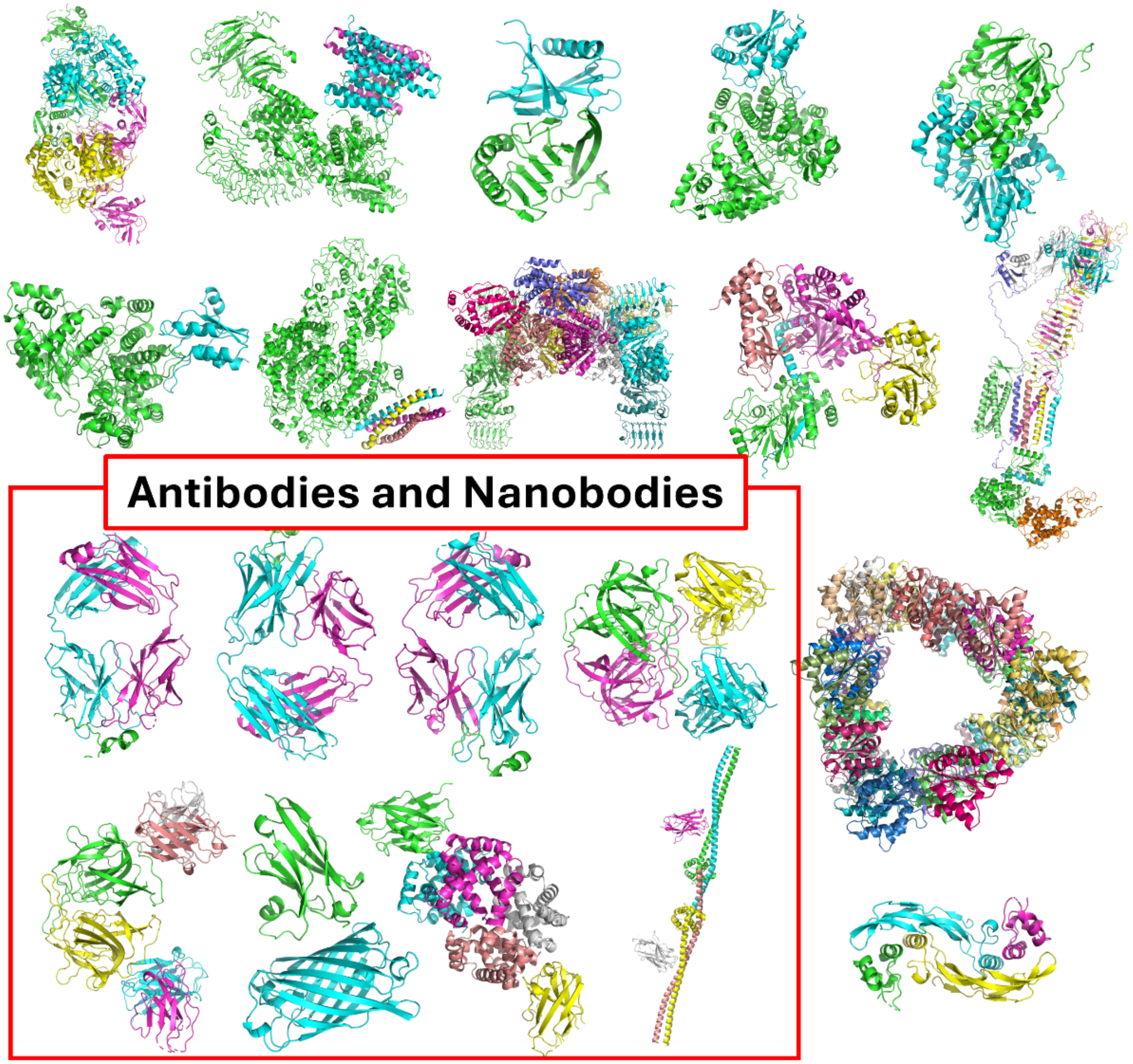


**Figure S1 (part 1)**. Overview of 40 assembly targets in CASP16.

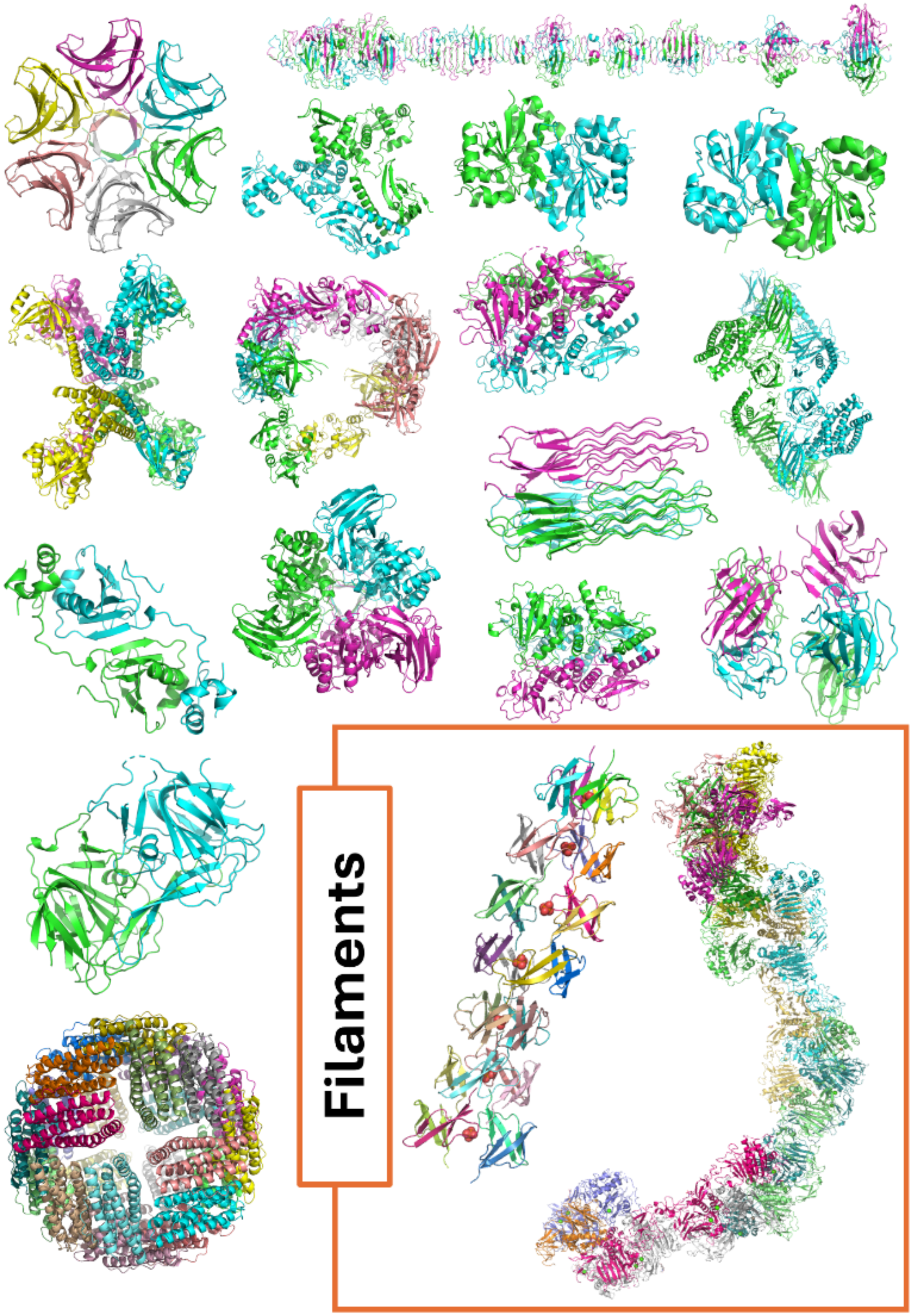


**Figure S1 (part 2)**. Overview of 40 assembly targets in CASP16.

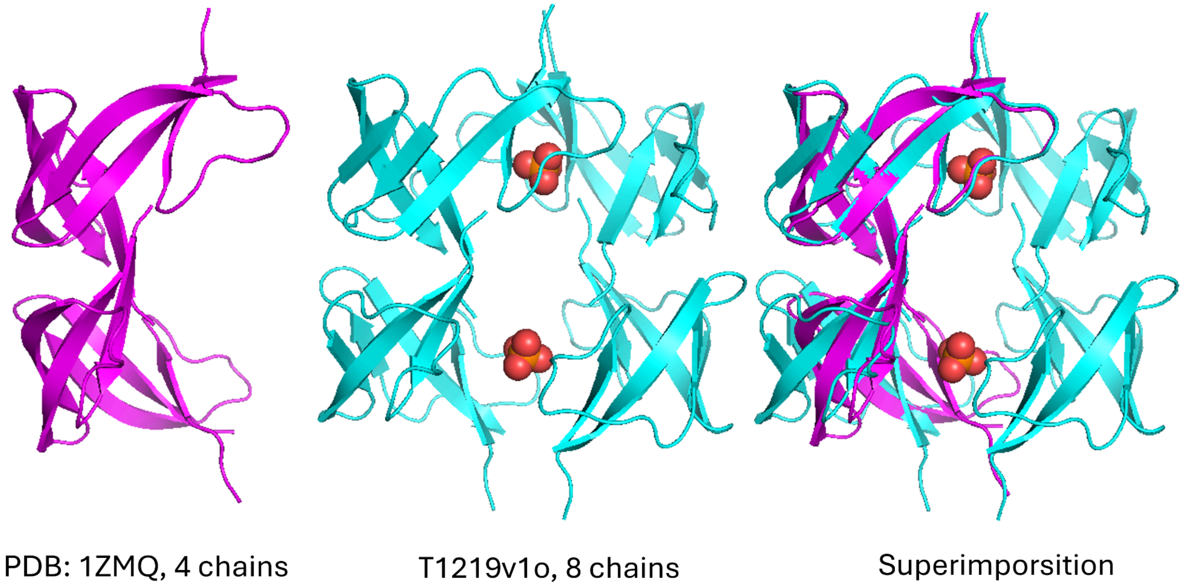


**Figure S2.** The comparison of 1ZMQ and current target (T1210v1o) structure. Both of them are the oligomers of human defensin-6. 1ZMQ is composed of 4 chains and solved by X-ray diffraction while the current target structure was solved by EM.

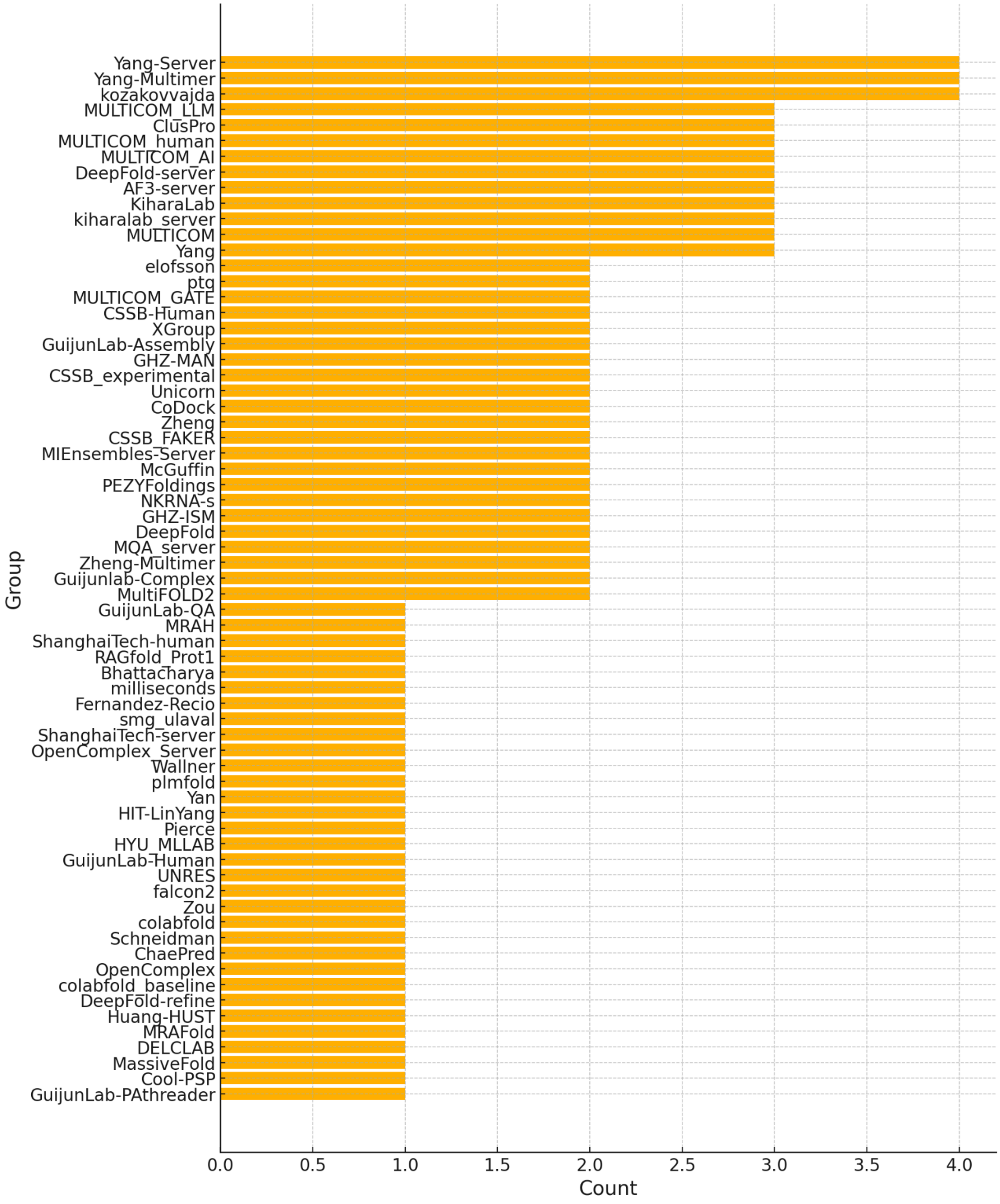


**Figure S3** The count of difficult targets for which each group was able to predict models with DockQ ≥ 0.5. Groups were excluded if they failed to produce any models with DockQ ≥ 0.5 on difficult targets. The 13 difficult targets were defined based on Figure 2A and include those shown in Figures 2B–I as well as antibody/nanobody–antigen assemblies.

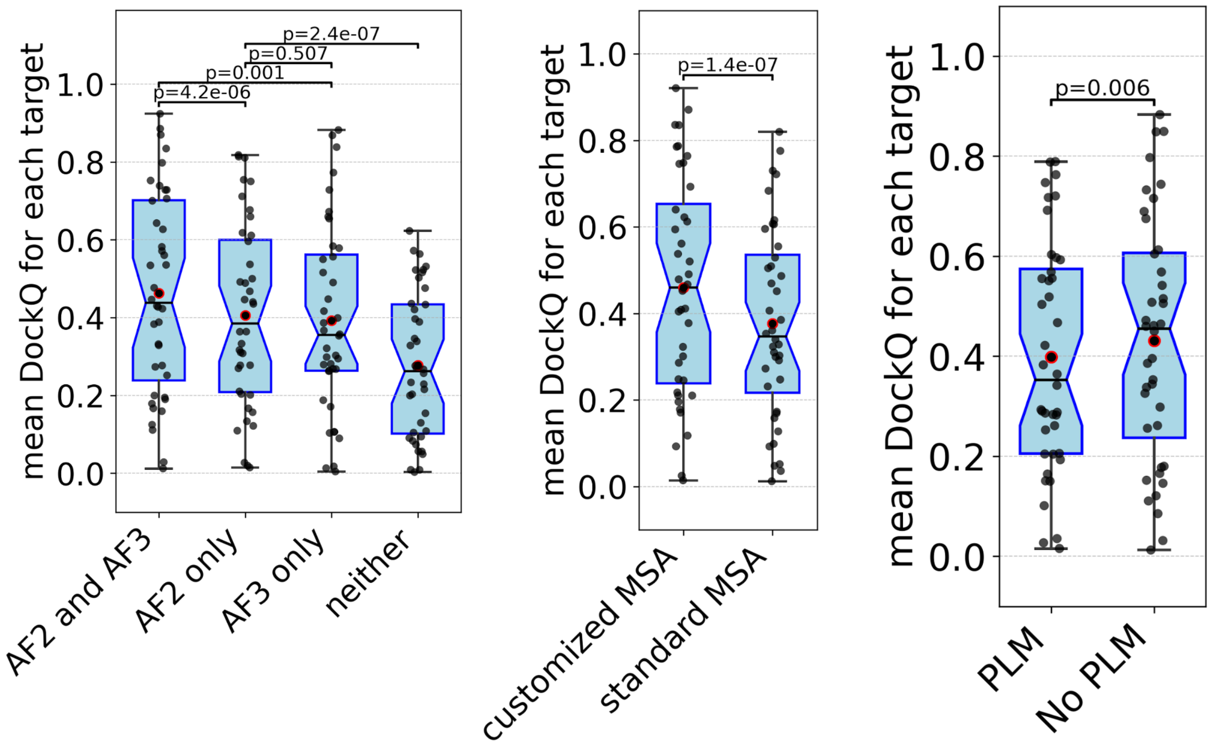


**Figure S4.** Relationship between methodologic features and prediction performance. Left: Groups categorized by use of AlphaFold2 (AF2) and/or AlphaFold3 (AF3). Middle: Groups using customized MSAs versus standard MSAs; Right: Groups using protein language models (PLMs) versus those that did not. For each target, the mean DockQ score of submitted models is shown. Groups incorporating both AF2 and AF3, customized MSAs, or PLMs tend to show higher performance.

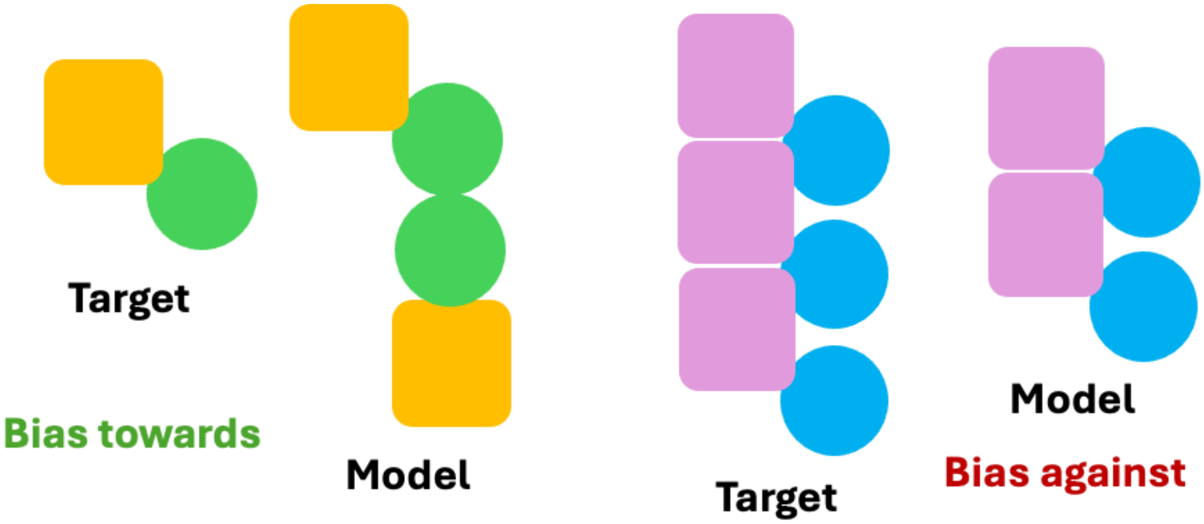


**Figure S5.** Scenarios where the OST pipeline will have undesirable bias. Left: a model with two incorrectly predicted interfaces, which will not be punished; right: a model with all the necessary interfaces to assemble into the target structure and no incorrectly predicted interfaces, which will be punished by OST due to missing chains appearing in the target.

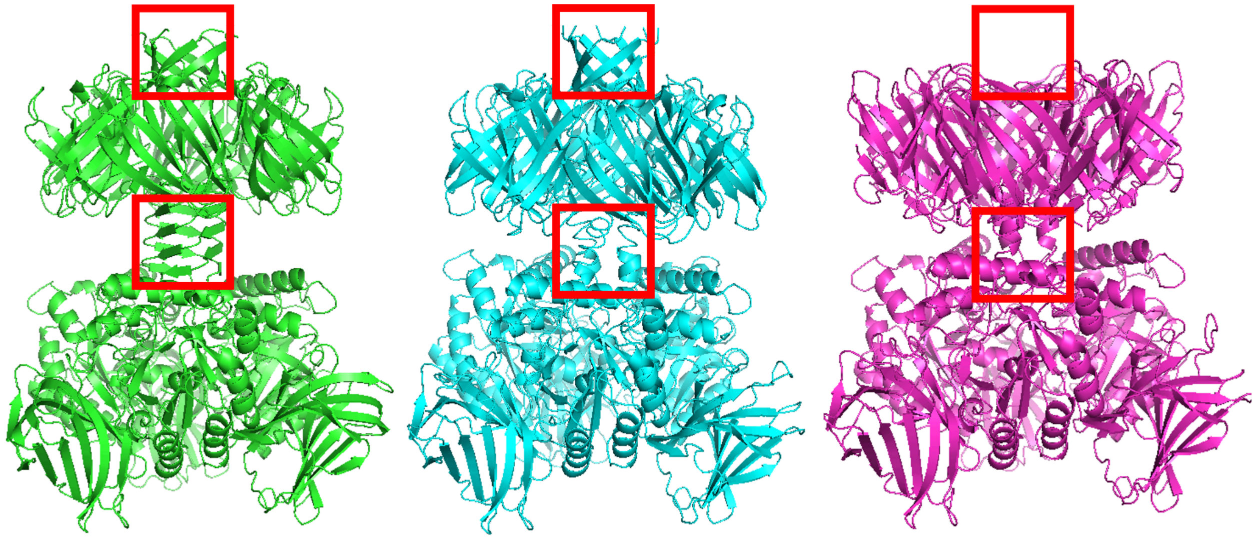

**Figure S6**. Structure of target H1236 and top-performing models. H1236 is a complex composed of six copies of gp30 and three copies of prokaryotic prokaryotic polysaccharide deacetylaseShown are the experimental structure of H1236 (left), the best model from KiharaLab (middle, DockQ = 0.615), and the best model from MassiveFold (right, DockQ = 0.379), the second-best performing group on this target. The red rectangular indicated regions participants usually failed to predict correct secondary structures and interactions.

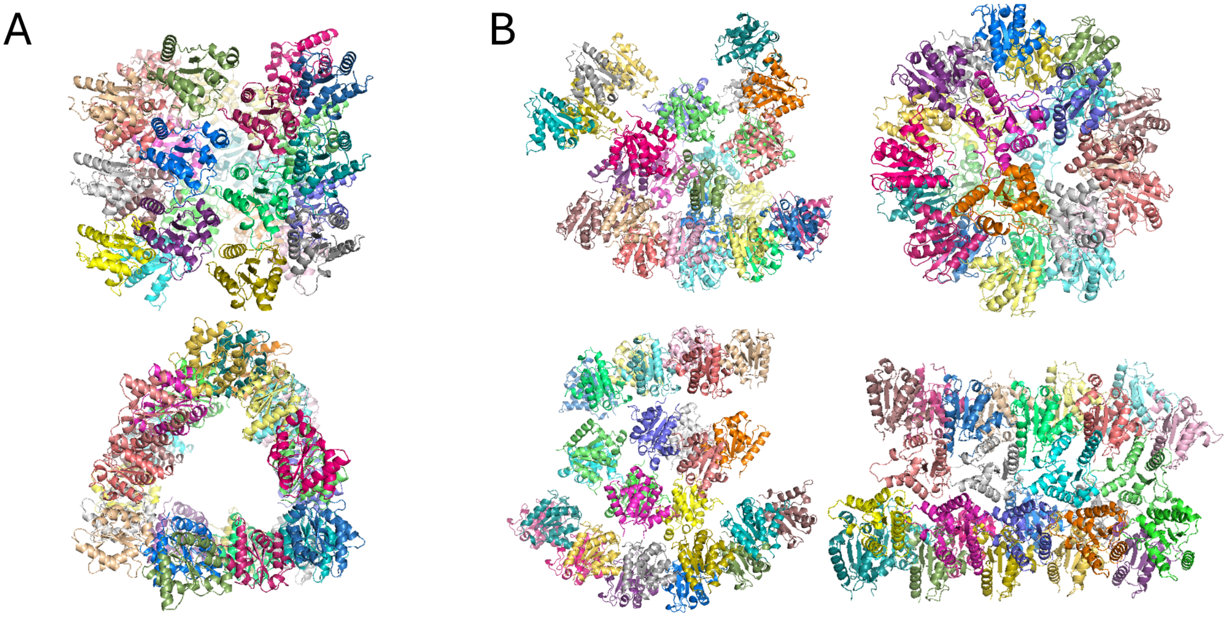


**Figure S7.** Structure of target H1265 and representative submitted models. (A) Experimental structure of H1265, a complex composed of TIR domains from TLR4 and MAL, shown from the side (top) and top-down (bottom) views. (B) Selected submitted models with either the relatively high TM-score, DockQ, or distinct alternative shape.

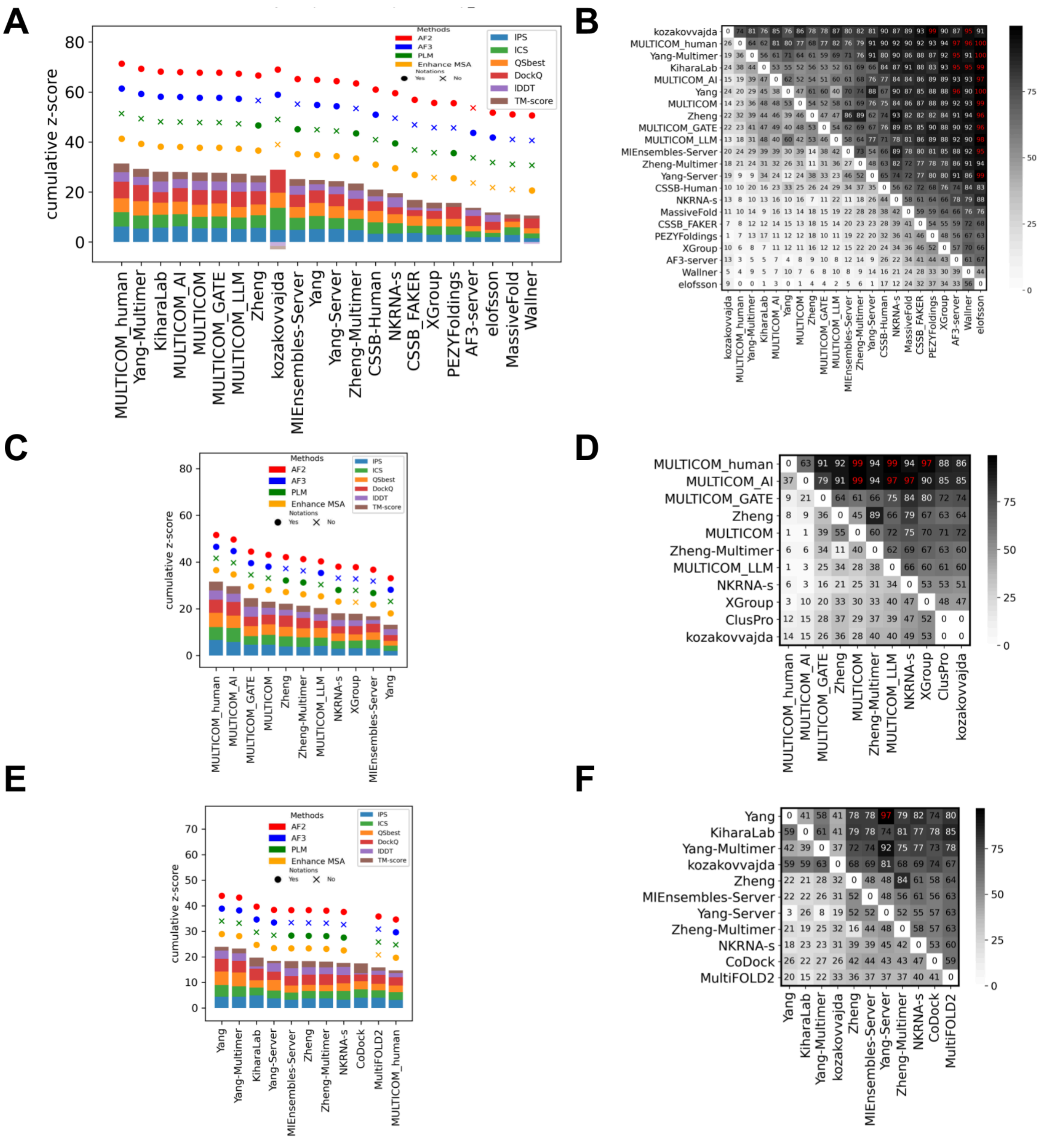


**Figure S8.** Rankings and head-to-head bootstrap comparison of top groups based on the first model for each group in phase 1 (A and B), phase 0 (C and D) and phase 2 (E and F).

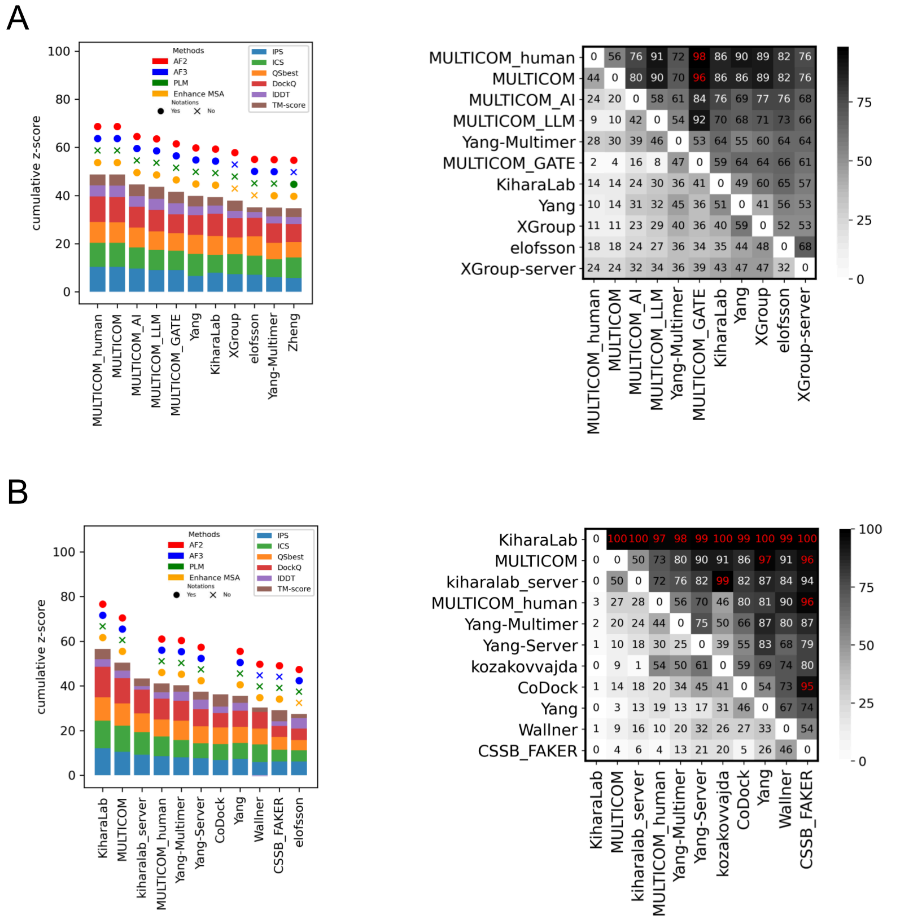


**Figure S9.** Rankings (left) and head-to-head bootstrap comparison (right) of top 11 groups based on the best model per group in phase 0 (A) and phase 2 (B).
